# Structural ordering of the *Plasmodium berghei* circumsporozoite protein repeats by inhibitory antibody 3D11

**DOI:** 10.1101/2020.06.02.131110

**Authors:** Iga Kucharska, Elaine Thai, Ananya Srivastava, John Rubinstein, Régis Pomès, Jean-Philippe Julien

## Abstract

*Plasmodium* sporozoites express circumsporozoite protein (CSP) on their surface, an essential protein that contains central repeating motifs. Antibodies targeting this region can neutralize infection, and the partial efficacy of RTS,S/AS01 – the leading malaria vaccine against *P. falciparum* (Pf) – has been associated with the humoral response against the repeats. Although structural details of antibody recognition of PfCSP have recently emerged, the molecular basis of antibody-mediated inhibition of other *Plasmodium* species via CSP binding remains unclear. Here, we analyze the structure and molecular interactions of potent monoclonal antibody (mAb) 3D11 binding to *P. berghei* CSP (PbCSP) using molecular dynamics simulations, X-ray crystallography, and cryoEM. We reveal that mAb 3D11 can accommodate all subtle variances of the PbCSP repeating motifs, and, upon binding, induces structural ordering of PbCSP through homotypic interactions. Together, our findings uncover common mechanisms of antibody evolution in mammals against the CSP repeats of *Plasmodium* sporozoites.

## INTRODUCTION

Despite extensive biomedical and public health measures, malaria persists as a major global health concern, with an estimated 405,000 deaths and 228 million cases annually (1). Moreover, resistant strains have been detected against all currently available antimalarial drugs, including sulfadoxine/pyrimethamine, mefloquine, halofantrine, quinine and artemisinin (2, 3). Although ~94% deaths are caused by *Plasmodium falciparum* (Pf) (1), other *Plasmodium* species that infect humans (*P. vivax, P. malariae, P. knowlesi* and *P. ovale*) also cause debilitating disease and have been associated with fatal outcomes (4). All *Plasmodium* species have a complex life cycle divided between a vertebrate host and an *Anopheles* mosquito vector (5). During a blood meal, sporozoites are deposited into the skin of a host organism from the salivary glands of a mosquito, and subsequently migrate through the bloodstream to infect host hepatocytes (6). Due to the small number of parasites transmitted and the expression of protein antigens that possess conserved functional regions (7, 8), the pre-erythrocytic sporozoite stage of the *Plasmodium* life cycle has long been considered a promising target for the development of an anti-malarial vaccine (9).

Circumsporozoite protein (CSP) is the most abundant protein on the surface of *Plasmodium* sporozoites, and is necessary for parasite development in mosquitoes and establishment of infection in host liver cells (10–12). Flanked by N- and C-terminal domains, CSP contains an unusual central region consisting of multiple, short (4 to 8) amino acid (aa) repeats (13–16). The sequence of the repeating motif depends on the *Plasmodium* species and field isolate (17–19). Importantly, the central region of CSP is highly immunodominant and antibodies targeting the repeats can inhibit sporozoite infectivity by preventing parasite migration (20) and attachment to hepatocytes (21, 22). PfCSP is a major component of the leading malaria vaccine RTS,S/AS01, which is currently undergoing pilot implementation in Africa (23, 24). Anti-PfCSP repeat antibodies have been suggested to form the predominant humoral immune response elicited by RTS,S/AS01, and they correlate with vaccine efficacy (25–27). However, RTS,S/AS01 offers only modest and short-lived protection (28–30); thus it is critical to develop a better molecular understanding of the antibody response against this *Plasmodium* antigen, particularly the repeat region (31–33), to provide valuable information needed for improved vaccine design.

Our understanding of *Plasmodium* biology and key host-parasite interactions has been enhanced by studies using rodent parasites, including *P. berghei* (Pb), *P. chabaudi* and *P. yoelii* (34). *In vivo* studies evaluating the inhibitory potential of mAbs are often derived from these rodent parasite models, or transgenic rodent sporozoites harboring PfCSP, as Pf fails to infect rodents. For example, mAb 3D11 was isolated from mice exposed to the bites of γ-irradiated Pb-infected mosquitoes (22). mAb 3D11 recognition of the central repeat of PbCSP on the surface of live sporozoites results in abolished Pb infectivity *in vitro* and *in vivo* (35). Electron micrographs of Pb sporozoites pre-treated with mAb 3D11 revealed the presence of amorphous, precipitated material on the parasite surface caused by the circumsporozoite precipitation reaction (22). This antibody continues to be widely used in model systems of sporozoite infection. For example, a recent study used mAb 3D11 in combination with transmission-blocking mAb 4B7 to show that antibody targeting of both the pre-erythrocytic and sexual stages of a Pfs25-transgenic Pb parasite led to a synergistic reduction of parasite transmission in mice (36). However, it remains unclear whether murine mAb 3D11 recognizes the central domain of PbCSP with the same molecular principles as the most potent human anti-PfCSP repeat antibodies, for which molecular details have recently emerged (37–44).

Here, we characterized the structure of the PbCSP repeats unliganded and as recognized by mAb 3D11. Our molecular studies reveal that mAb 3D11 binds across all PbCSP repeat motifs and induces structural ordering of PbCSP in a spiral-like conformation using homotypic interactions.

## RESULTS

### Repeat motifs of PfCSP and PbCSP have similar structural propensities

The central repeat of PfCSP and PbCSP consist of recurring 4-aa motifs rich in asparagine and proline residues (**Fig. 1A**). PfCSP is composed of NANP repeating motifs interspersed with intermittent NVDP repeats, and contains a singular NPDP motif in the junction immediately following the N-terminal domain. While the major repeat motif of PfCSP has often been referred to as NANP, numerous reports have identified NPNA as the structurally relevant unit of the central region (39, 42, 45, 46). Similarly, the central domain of PbCSP contains an array of PPPP and PAPP motifs interspersed with NPND or NAND motifs (**Fig. 1A**). Notably, both orthologs contain the conserved pentamer, KLKQP, known as Region I, at the C-terminal end of the N-terminal domain.

**Figure 1.**
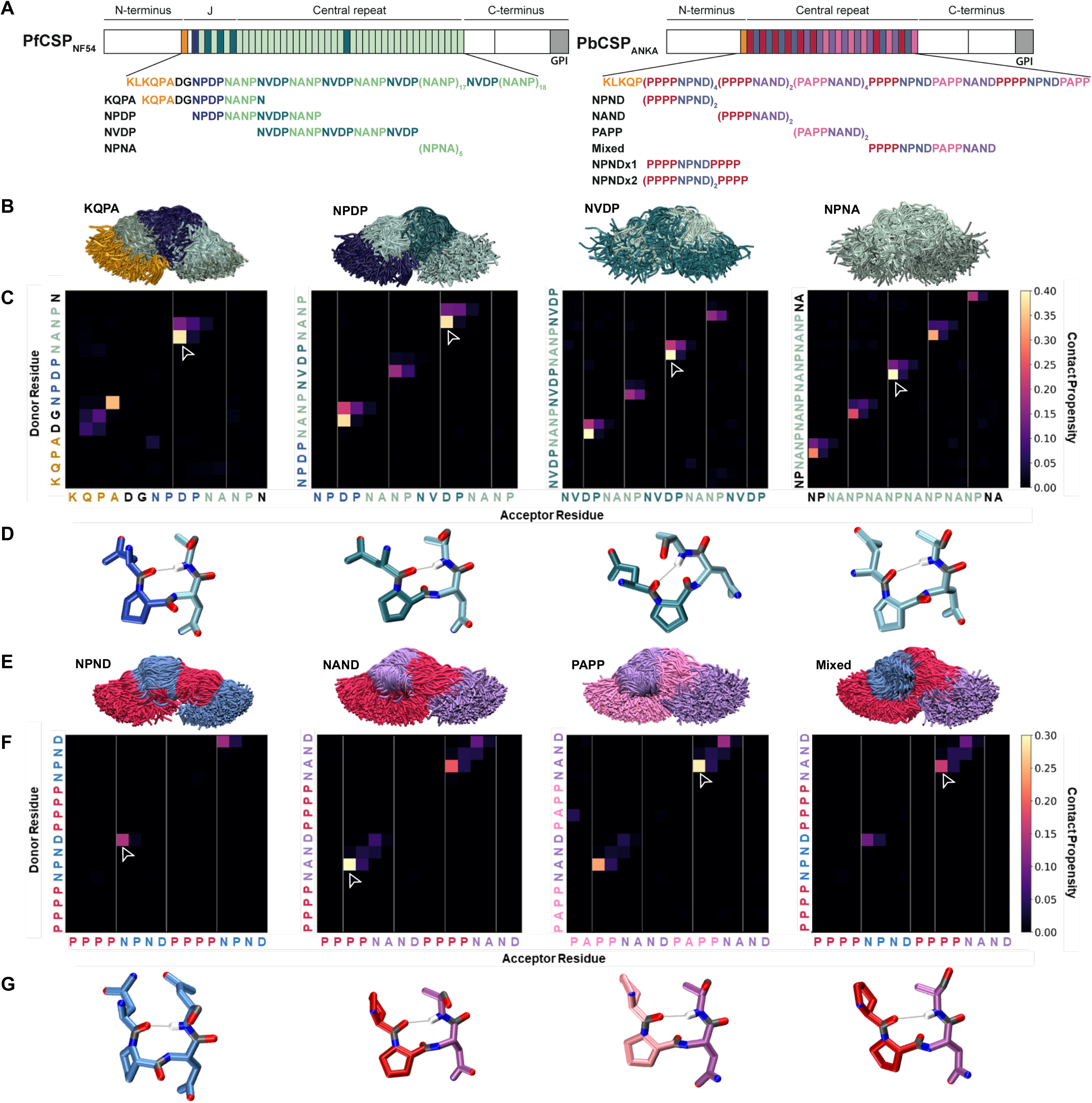
Comparison of PfCSP and PbCSP repeat sequences and structures. **(A)** Schematic representations of PfCSP strain NF54 and PbCSP strain ANKA, each comprising an N-terminal domain, central repeat region, and C-terminal domain. The junctional region (J) immediately following the N-terminal domain of PfCSP is indicated. Colored bars represent each repeat motif. The sequences of each CSP central repeat region and corresponding peptides used in the study are shown below their respective schematics. **(B-G)** Conformational ensembles of CSP peptides in solution from molecular dynamics simulations. **(B)** Superposition of the conformations of the four PfCSP-derived peptides at each nanosecond. The peptides are aligned to the conformational median structure and only the backbone is shown for clarity. **(C)** Ensemble-averaged backbone-backbone hydrogen-bonding maps for each PfCSP peptide sequence. The propensity for hydrogen bonds between the NH groups (y-axis) and CO groups (x-axis) is indicated by the color scale on the right. **(D)** Sample molecular dynamics snapshots of the highest-propensity turn for each PfCSP peptide are shown as sticks with hydrogen bonds shown as grey lines. The highest-propensity turn for each peptide is indicated by the arrowhead on the corresponding hydrogenbonding map. **(E)** Superposition of the conformations of the four PbCSP-derived peptides at each nanosecond. The peptides are aligned to the conformational median structure and only the backbone is shown for clarity. **(F)** Ensemble-averaged backbone-backbone hydrogen-bonding maps for each PbCSP peptide sequence. The propensity for hydrogen bonds between the NH groups (y-axis) and CO groups (x-axis) is indicated by the color scale on the right. **(G)** Sample molecular dynamics snapshots of the highest-propensity turn for each PbCSP peptide are shown as sticks with hydrogen bonds shown as grey lines. The highest-propensity turn for each peptide is indicated by the arrowhead on the corresponding hydrogenbonding map.

To examine and compare the structural properties of the various Pf and Pb repeat motifs in solution, we performed molecular dynamics (MD) simulations using eight different peptides ranging in length from 15-20 aa, with four peptides derived from Pf [KQPADGNPDPNANPN (“KQPA”); NPDPNANPNVDPNANP (“NPDP”); (NVDPNANP)_2_NVDP (“NVDP”); and (NPNA)_5_ (“NPNA”)], and four peptides from Pb [(PPPPNPND)_2_ (“NPND”); (PPPPNAND)_2_ (“NAND”); (PAPPNAND)_2_ (“PAPP”); and PPPPNPNDPAPPNANAD (“Mixed”); **Fig. 1B-G**]. Each simulation was conducted in water for a total production time of 18 μs. All eight peptides were highly disordered and adopted a large ensemble of conformations with low to moderate secondary structure propensities (**Fig. 1B** and **E**), which are best described in statistical terms. The only secondary structure observed was local, and consisted of sparse, transient hydrogen-bonded turns (**Figs. 1C** and **F**, **S1** and **S2**, and **Supplemental Table 1**). In particular, these interactions consisted of forward α-, β-, and γ-turns, with the β-turns being the most populated (up to 40%; **Fig. 1C** and **F**), consistent with previous NMR studies focused on the NANP repeats (45). Across all peptides, the average β-turn lifetime ranged from 2.7 ± 0.2 ns for the Pb PPNA turn to 4.4 ± 0.4 ns for DPNA turns found within PfCSP (**Supplemental Table 1**).

In line with reports identifying NPNA as the main structural repeat motif of PfCSP (39, 42, 45, 46), turns were predominantly observed within these motifs, as well as DPNA, NPNV and ADGN sequences amongst the PfCSP peptides. NPND and PPNA exhibited the greatest propensity to form β-turns of the PbCSP motifs. Importantly, each individual motif consistently exhibited the same structural tendencies, independent of their position and the overall peptide sequence in which they were contained (**Fig. 1C** and **F**, and **Supplemental Table 1**). Furthermore, using the probability rule stating that two events are independent if the equation P(A⋂B) = P(A)·P(B) holds true, we show that the presence of an intramolecular hydrogen bond in one motif does not alter the hydrogen-bonding propensities of adjacent motifs (**Fig. S2B** and **C**). Therefore, we conclude that there is no discernable cooperativity between the structures of the different repeat motifs, and as such, in the absence of extended or nonlocal secondary structure, each of these motifs behaves as an independent unit with its own intrinsic secondary structure propensities.

To examine the influence of Asn, Asp, and Gln sidechains on the conformational ensemble of the peptides, we computed contact maps for backbone-sidechain hydrogen bonds (**Fig. S1** and **Supplemental Table 1**). We found that the majority of contacts are in the form of pseudo α-turns and β-turns, with backbone NH groups donating to sidechain O atoms. Notably, we discovered that these transient sidechain contacts do not have a stabilizing effect on backbone-backbone hydrogen bonds and consequently, are not correlated with the presence of these bonds (numerical example in **Fig. S2B** and **C**).

In summary, the four Pf and four Pb peptides corresponding to CSP central repeats were all found to be highly disordered, resulting in an ensemble of conformations. The only secondary structure elements present in the ensemble of conformations were sparse and local hydrogen-bonded turns within each motif. Each structural motif acted independently from adjacent sequences and behaved similarly in various peptides.

### Multiple copies of mAb 3D11 bind PbCSP with high affinity

Next, we investigated the binding of mAb 3D11 to the PbCSP repeat of low structural propensity. Our biolayer interferometry (BLI) studies indicated that Fab 3D11 binds PbCSP with complex kinetics, but overall high affinity (**Fig. 2A**). Isothermal titration calorimetry (ITC) also indicated a high affinity interaction, with a K_D_ value of 159 ± 47 nM (**Fig. 2B**). In addition, ITC revealed a very high binding stoichiometry (N = 10 ± 1), suggesting that approximately ten copies of 3D11 Fab bound one molecule of PbCSP simultaneously. Size exclusion chromatography coupled with multi-angle light scattering (SEC-MALS) characterization of the 3D11 Fab-PbCSP complex confirmed the high binding stoichiometry with a molecular weight of 587 ± 7 kDa for the complex (**Fig. 2C-D**). This size is consistent with approximately eleven 3D11 Fabs bound to one molecule of PbCSP, and thus in agreement with the results from the ITC studies within experimental error. Therefore, through a number of biophysical studies, we show that up to eleven copies of 3D11 Fab can bind simultaneously to PbCSP with high affinity.

**Figure 2.**
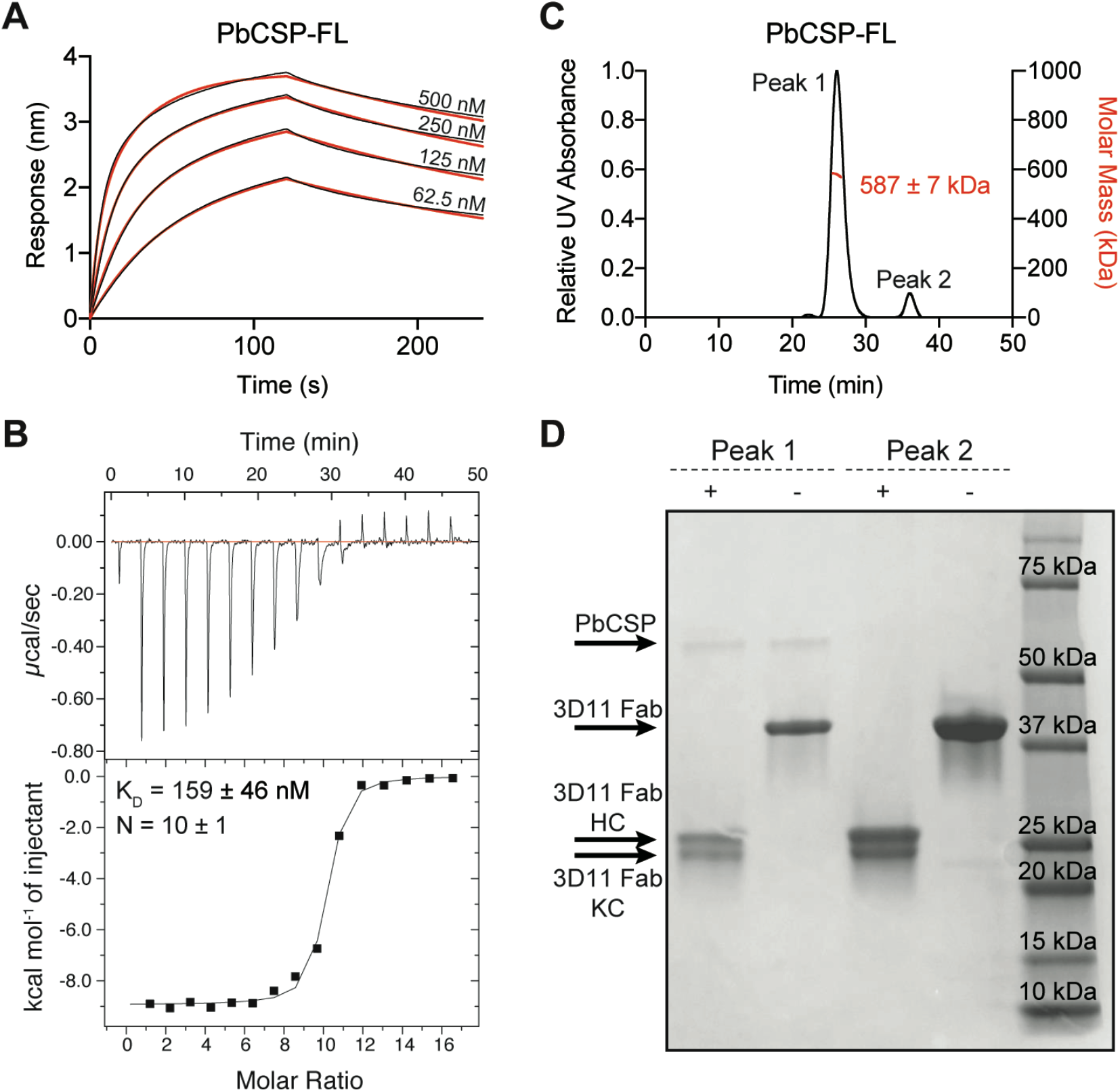
Biophysical characterization of 3D11 Fab-PbCSP binding. **(A)** Binding kinetics of twofold dilutions of 3D11 Fab to full-length PbCSP. Representative sensorgrams are shown in black and 2:1 model best fits in red. Data are representative of three independent measurements. **(B)** Isothermal titration calorimetry (ITC) analysis of 3D11 Fab binding to full-length PfCSP at 37°C. Above, raw data of 16 injections of 3D11 Fab (0.2 mM) into the sample cell containing PbCSP (0.02 mM). Below, plot and trendline of heat of injectant corresponding to the raw data. K_D_ and N values resulting from three independent experiments are indicated. Standard error values are reported as standard error of the mean (SEM). **(C)** Results from size exclusion chromatography coupled with multi-angle light scattering (SEC-MALS) for the 3D11 Fab-PbCSP complex. A representative measurement of the molar mass of the 3D11 Fab-PbCSP complex is shown as the red line. Mean molar mass and SD are as indicated. **(D)** SDS-PAGE analysis of resulting Peaks 1 and 2 from SEC-MALS. Each peak was sampled in reducing and non-reducing conditions as indicated by + and −, respectively.

### mAb 3D11 is cross-reactive with subtly different PbCSP motifs in the central repeat

We next sought to define the exact mAb 3D11 epitope. We first conducted BLI studies to confirm that mAb 3D11 does not bind the PbCSP C-terminal domain (residues 202-318; **Fig. S3A**). Next, we performed ITC studies to evaluate 3D11 Fab binding to each of the four peptides derived from the PbCSP central repeat region that were used in our MD simulations (**Fig. 3A**). Our experiments revealed that mAb 3D11 preferentially binds the NPND and Mixed peptides with high affinity (K_D_ = 45 ± 15 nM and 44 ± 4 nM, respectively), but also binds the NAND and PAPP peptides, albeit with lower affinity (K_D_ = 207 ± 1 nM and 611 ± 139 nM, respectively).

**Figure 3.**
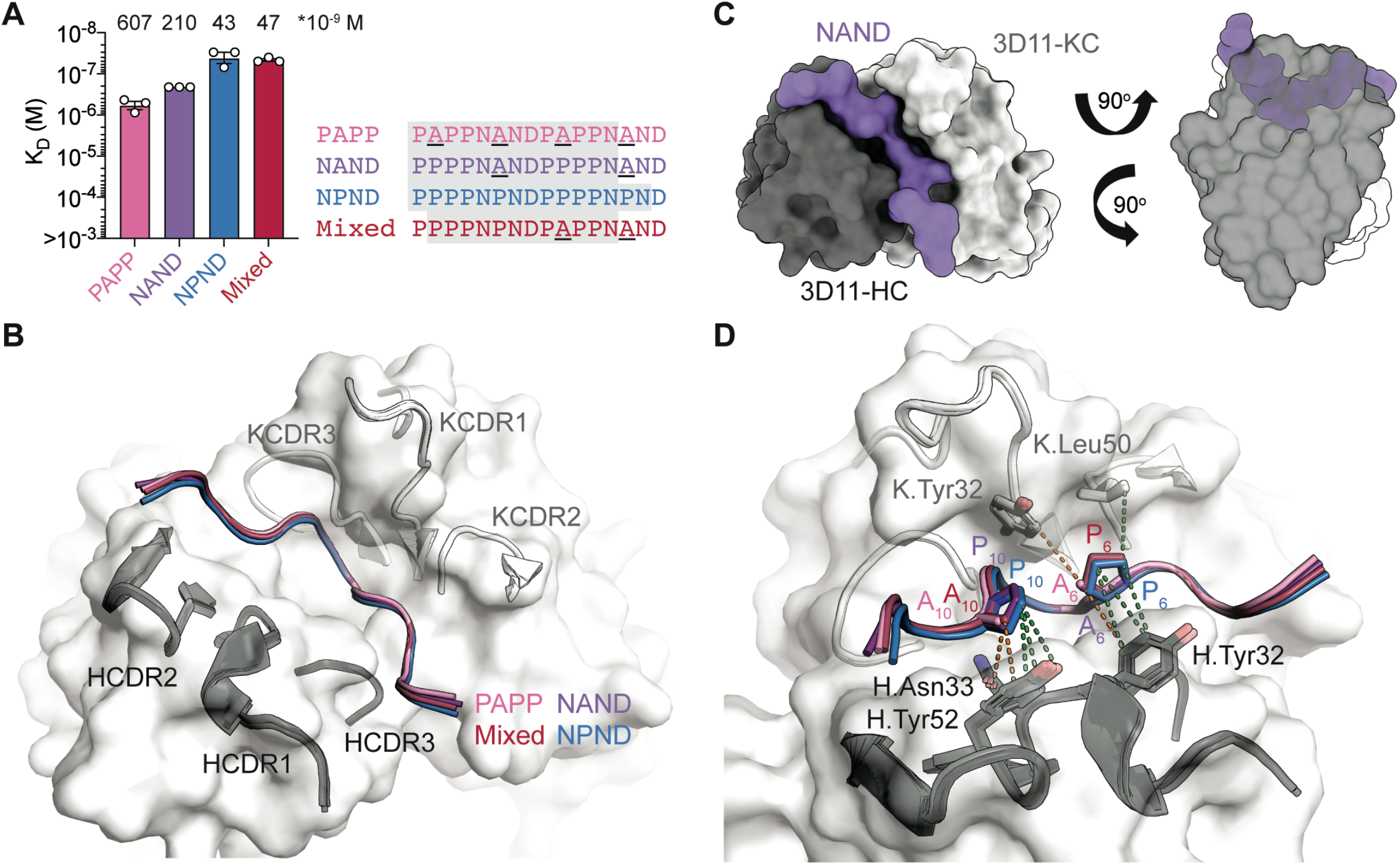
3D11 Fab binding to PbCSP repeat peptides. **(A)** Affinities of 3D11 Fab for PAPP, NAND, NPND and Mixed peptides as measured by ITC. Symbols represent independent measurements. Mean K_D_ values are shown above the corresponding bar. Error bars represent SEM. **(B)** The 3D11 Fab binds the PAPP (pink), NAND (purple), NPND (blue) and Mixed (red) peptides in nearly identical conformations. mAb 3D11 CDRs are indicated. **(C)** Overview and side view of the NAND peptide (purple) in the binding groove of the 3D11 Fab shown as surface representation (H-chain shown in black and K-chain shown in grey). **(D)** Van der Waals interactions formed by side chain atoms of both Ala and Pro residues are indicated by orange dashed lines, and those unique to Pro6 and Pro10 are indicated by green dashed lines.

To gain insight into the molecular basis of this preference, we solved the X-ray crystal structures of 3D11 Fab in complex with each peptide. The structure of the 3D11 Fab-NPND complex was determined at 2.30 Å resolution, while the structures of 3D11 Fab in complex with each of the other three peptides were all solved at ~1.60 Å resolution (**Table 1**). Interestingly, all four peptides adopted almost identical conformations when bound by 3D11 Fab (**Fig. 3B** and **Fig. S3**), fitting deep into the binding groove and forming a curved U-shape structure (**Fig. 3C**). Amongst all four peptides, the mAb 3D11 core epitope consisted of eight residues [PN(A/P)NDP(A/P)P] with an all-atom RMSD < 0.5 Å. Importantly, this shared recognition mode ideally positions aromatic side chains in the mAb 3D11 complementarity determining regions (CDRs) to form favorable pi-stacking and hydrophobic cage interactions around each PbCSP peptide (**Fig. S4**). Indeed, the majority of these contacts are made with residues that are conserved between all PbCSP repeat motifs, and thus contribute to the cross-reactive binding profile of mAb 3D11.

**Table 1:**
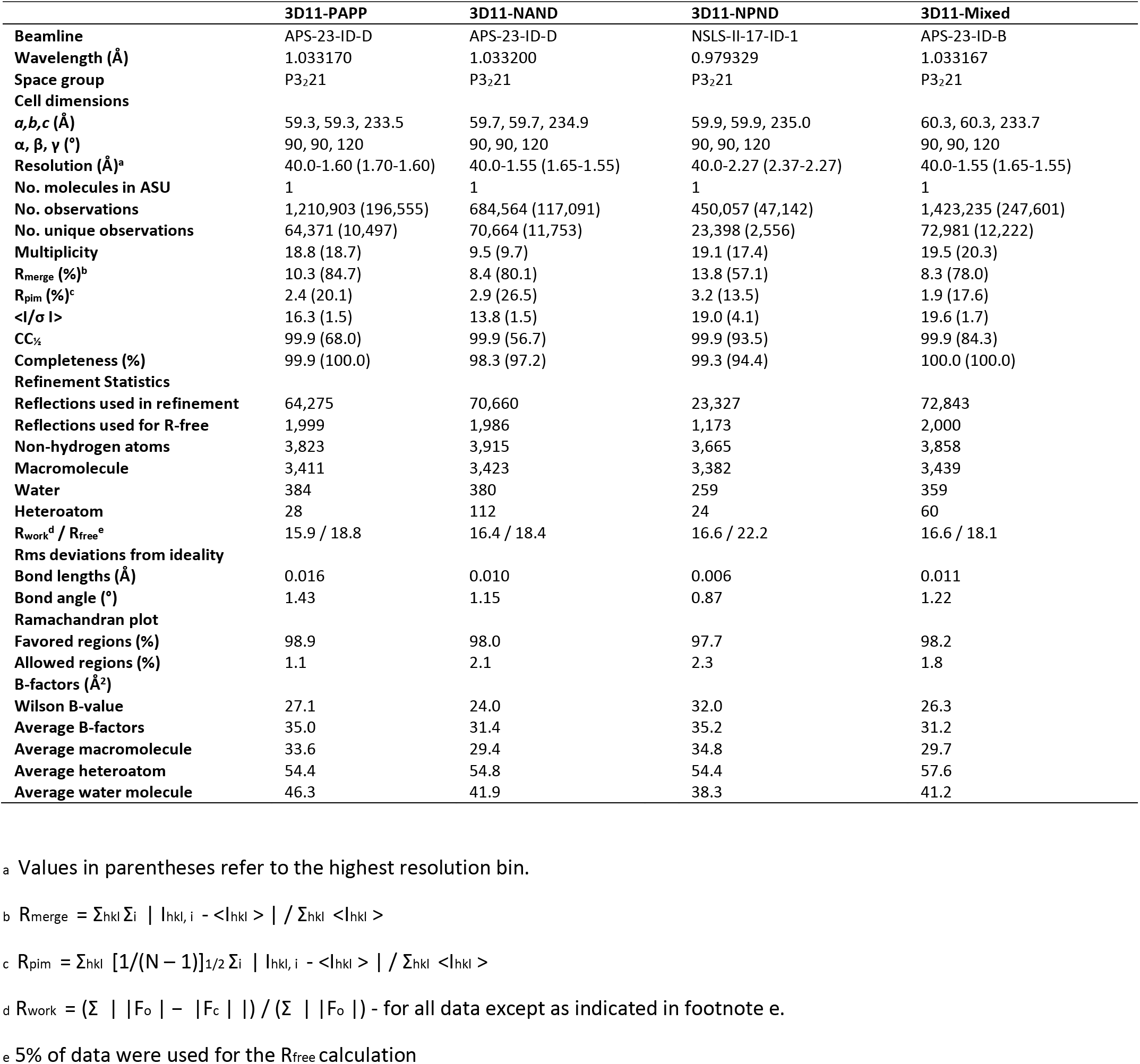
X-ray crystallography data collection and refinement statistics.

Despite binding in nearly identical conformations, differences exist in the molecular details of 3D11 Fab binding to each peptide that provide key insights into mAb 3D11 recognition of PbCSP. Our crystal structures revealed that more van der Waals contacts were formed by a Pro residue in the PPPP and NPND motifs compared to an Ala at the same position in the PAPP and NAND motifs (**Fig. 3D**). Consequently, the epitopes of the NAND, NPND and Mixed peptides had a slightly greater buried surface area (BSA; 753, 762, and 765 Å^2^, respectively) as compared to the PAPP peptide (743 Å^2^), which only consists of Ala-containing motifs (**Supplemental Table 2**). In particular, Pro10 of the PPPP motif found in the NAND and NPND peptides forms more van der Waals interactions with antibody residues H.Asn33 and H.Tyr52 compared to Ala10 of the PAPP motif present in PAPP and Mixed peptides. Similarly, Pro6 of the NPND motif in the NPND and Mixed peptides makes additional interactions with antibody residue K.Leu50 that are not present for Ala6 of the NAND motif within the PAPP and NAND peptides (**Supplemental Table 2**). These differences in interactions observed at the atomic level directly relate to the binding affinities measured by ITC, where the PbCSP peptides that bury more surface area in the 3D11 paratope show the highest binding affinities (**Fig. 3A**).

### 3D11 binding stabilizes the central PbCSP repeat in a spiral-like conformation

To understand how mAb 3D11 recognizes full-length PbCSP, we performed cryoEM analysis of the SEC-purified 3D11 Fab-PbCSP complex (**Fig. 2D**). A dataset of 165,747 3D11 Fab-PbCSP particle images was refined with no symmetry imposed, resulting in a 3.2 Å resolution reconstruction showing seven predominant 3D11 Fabs peripherally arranged around PbCSP with their variable domains clustered around a central density (**Fig. 4**, **Table 2** and **Fig. S5**). The PbCSP repeat forms the core of the complex and is arranged into a triangular spiral of 51 Å pitch and 16 Å diameter (**Fig. 4**), which fits 61 out of 108 residues of the PbCSP central region. We assigned the density to the high-affinity PPPPNPND repeats.

**Figure 4.**
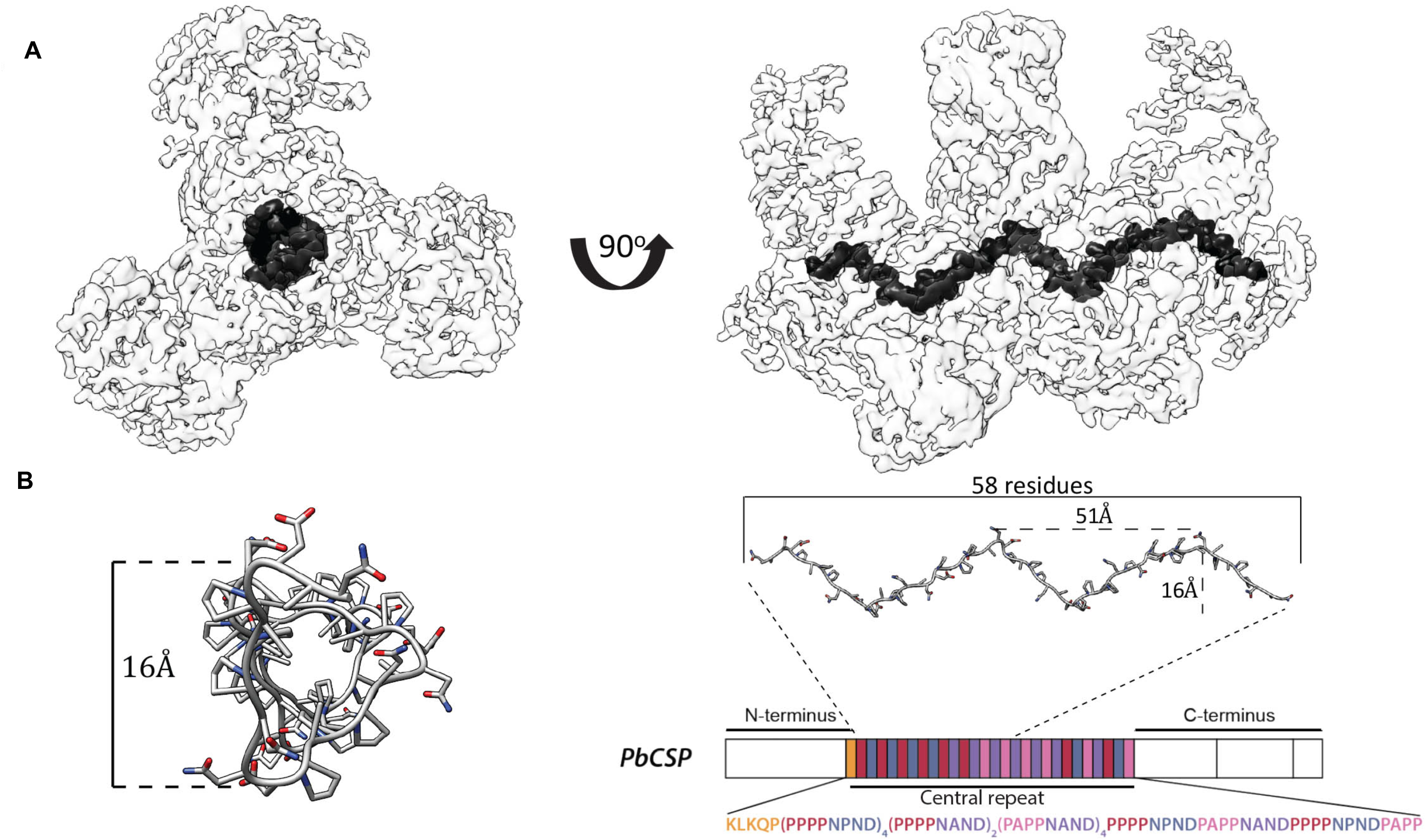
Spiral organization of the PbCSP repeat upon 3D11 Fab binding. **(A)** CryoEM map of the 3D11 Fab-PbCSP complex is shown as a transparent light gray surface with the PbCSP density highlighted in black. **(B)** The PbCSP model built into the cryoEM map is shown in dark grey as sticks and aligned to the schematic representation of the PbCSP protein sequence.

**Table 2.** CryoEM data collection and refinement statistics.

| **Data Collection** |  |
| --- | --- |
| **Electron microscope** | Titan Krios G3 |
| **Camera** | Falcon 3EC |
| **Voltage (kV)** | 300 |
| **Nominal magnification** | 75,000 |
| **Calibrated physical pixel size (Å)** | 1.06 |
| **Total exposure (e<sup>-</sup> / Å<sup>2</sup>)** | 42.7 |
| **Number of frames** | 30 |
| **Image Processing** |  |
| **Motion correction software** | cryoSPARCv2 |
| **CTF estimation software** | cryoSPARCv2 |
| **Particle selection software** | cryoSPARCv2 |
| **3D map classification and refinement software** | cryoSPARCv2 |
| **Micrographs used** | 2,080 |
| **Particles selected** | 669,223 |
| **Global resolution (Å)** | 3.2 |
| **Particles contributing to final maps** | 165,747 |
| **Model Building** |  |
| **Modeling software** | Coot, phenix.real_space_refine |
| **Number of residues built** | 3,085 |
| **RMS (bonds)** | 0.002 |
| **RMS (angles)** | 0.56 |
| **Ramachandran favored (%)** | 95.8 |
| **Rotamer outliers (%)** | 0.5 |
| **Clashscore** | 6.27 |
| **MolProbity score** | 1.63 |
| **EMRinger score** | 2.54 |

The angle between two Fab variable domains is ~126°, such that approximately three Fabs are required to complete one full turn of the spiral (**Fig. 5A-B**). The cryoEM structure of the 3D11 Fab-PbCSP complex and the crystal structures of the 3D11 Fab-peptide complexes are in remarkable agreement for both the Fab (RMSD=0.39 Å) and the PbCSP repeat region (RMSD=0.66 Å) (**Fig. S6**). Minor differences exist in the N and C termini of the peptides, presumably because the termini are largely unrestricted in the crystal structure compared to the cryoEM structure.

**Figure 5.**
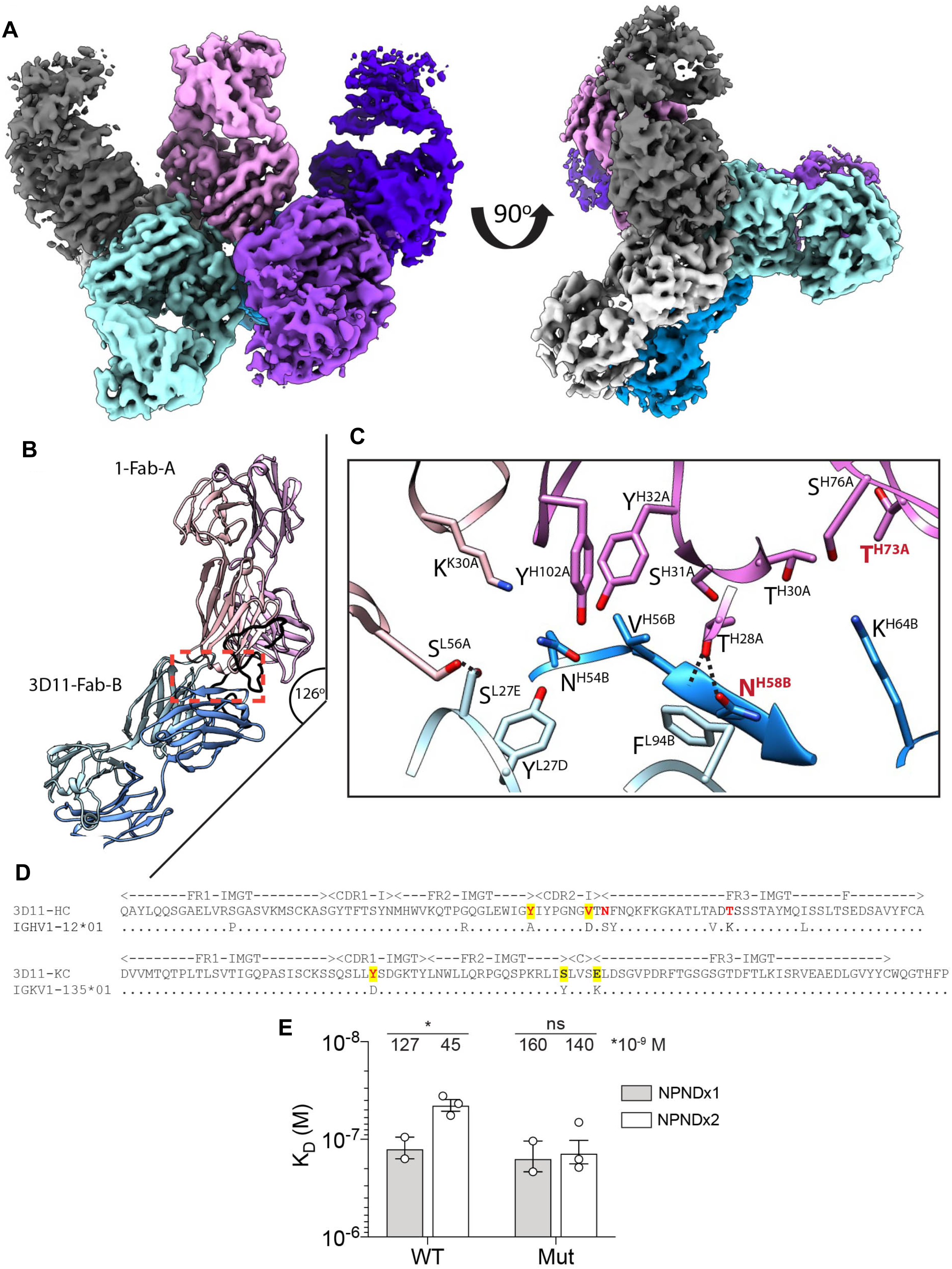
Homotypic interactions between 3D11 Fabs stabilize the 3D11 Fab-PbCSP complex. **(A)** The cryoEM map of the 3D11 Fab-PbCSP complex contains density for seven predominant 3D11 Fabs. Densities of individual Fabs are colored from pink to grey. **(B)** Close-up view of two adjacent 3D11 Fabs (pink and blue) from the cryoEM structure in complex with PbCSP (black). A red box denotes the site of Fab-PbCSP and Fab-Fab interactions. **(C)** Details of 3D11 Fab-Fab homotypic interactions. Residues forming Fab-Fab contacts are labelled in black. mAb 3D11 affinity matured residues that engage in Fab-Fab contacts but do not directly interact with PbCSP are labelled in red. Black dashes indicate H-bonds between residues. **(D)** Sequence alignment of mAb 3D11 with inferred germline precursors for the heavy and light chains (upper and lower row, respectively). Affinity matured residues that contact PbCSP are highlighted in yellow and affinity matured residues involved in homotypic interactions are indicated in red. **(E)** Binding affinity of WT 3D11 and H-58/73 germline-reverted mutant (Mut) Fabs to NPNDx1 (grey bars) and NPNDx2 (white bars) peptides as measured by ITC. Symbols represent independent measurements. Mean K_D_ values resulting from at least two independent experiments are shown. Error bars represent standard error of the mean. An unpaired one-tailed t-test was performed using GraphPad Prism 8 to evaluate statistical significance: *P < 0.05.

### Contacts between 3D11 Fabs stabilize the PbCSP spiral structure

To access their repeating and densely-packed epitopes, 3D11 Fabs are closely arranged against one another in the 3D11 Fab-PbCSP complex. Indeed, the epitope for a single Fab can be defined by 14 residues (PPPPNPNDPPPPNP, **Supplemental Table 3**), with the six C-terminal residues constituting the beginning of the epitope for the adjacent Fab. When considering two adjacent Fabs as a single binding unit, the BSA of the Fabs is 1313 Å^2^, and 1636 Å^2^ for PbCSP. Interestingly, we observe multiple Fab-Fab contacts in the cryoEM structure (**Fig. 5C** and **Fig. S7**). Comparison of the mAb 3D11 sequence to its inferred germline precursor (IGHV1-12 and IGKV1-135) reveals that some of the residues involved in homotypic contacts have been somatically hypermutated (H.Tyr50, H.Val56, H.Asn58, H.Thr73 and K.Tyr27D; **Fig. 5C-D**). In contrast to H.Tyr50, H.Val56 and K.Tyr27D that mediate Fab-Fab contacts as well as interact directly with PbCSP, H.Asn58 and H.Thr73 are only involved in Fab-Fab interactions without contacting PbCSP.

To investigate the role of affinity maturation in enhancing Fab-Fab contacts, somatically mutated heavy chain (HC) residues H.Asn58 and H.Thr73 were reverted to their inferred germline precursors (N58S and T73K: subsequently named H-58/73; **Fig. 5D**). We performed ITC studies to evaluate binding of wild-type (WT) and H-58/73 germline-reverted mutant 3D11 Fabs to two peptides derived from PbCSP, designed based on our X-ray and cryoEM structures to constitute the minimal binding site for one 3D11 Fab (PPPPNPNDPPPP, denoted “NPNDx1”) or two 3D11 Fabs in a “head-to-head” conformation (PPPPNPNDPPPPNPNDPPPPNPND, denoted “NPNDx2”). Although both WT and H-58/73 germline-reverted mutant Fabs bound NPNDx1 with comparable affinity, WT 3D11 Fab demonstrated significantly greater affinity for NPNDx2 compared to NPNDx1 (KD values of 45 ± 6 nM and 127 ± 32 nM, respectively; **Fig. 5E**). On the other hand, the H-58/73 germline-reverted mutant bound each peptide with similar affinities (KD values of 140 ± 38 nM for NPNDx2 and 160 ± 56 nM for NPNDx1; **Fig. 5E**). The improved binding affinity of WT 3D11 Fab for NPNDx2 compared to NPNDx1 which is not observed for the H-58/73 germline-reverted mutant 3D11 Fab suggests an important role for residues H.Asn58 and H.Thr73 in mediating homotypic interactions between neighboring 3D11 Fabs bound to PbCSP. Together, these data provide evidence for the affinity maturation of homotypic contacts that indirectly strengthen mAb 3D11 affinity to PbCSP.

## DISCUSSION

The CSP repeat is of broad interest for malaria vaccine design because it is targeted by inhibitory antibodies capable of preventing sporozoite infection as the parasite transits from *Anopheles* mosquitoes to mammalian hosts. Biophysical studies of the PfCSP central NANP repeat have shown that this region possesses low secondary structure propensities (45, 47–49), and AFM studies on live Pf sporozoites suggest a range of conformations for PfCSP (50, 51). Recent studies have uncovered that some of the most potent antibodies at inhibiting sporozoites are crossreactive with the PfCSP N-junction region that harbors KQPA, NPDP and NVDP motifs interspersed with NANP motifs (39, 40, 43). Our MD simulations of different sub-regions of the PfCSP central repeat including the N-junction provided detailed descriptions of their conformational ensemble and revealed that all sequences possess similar low structural propensity.

Our MD simulations for PbCSP also indicated that the low structural propensity of central repeat motifs with subtle sequence variance extends to other *Plasmodium* species. These findings are in agreement with studies linking repetitive, low-complexity peptide sequences to structural disorder (47–49). The role of the numerous repetitive sequences observed in parasitic genomes (52–54) remains to be fully understood, but is postulated to include maximizing parasite interactions with the target host cell (53), allowing the parasite to adapt under selective pressure by varying its number of repeats (55), and impairing the host immune response (56–58).

Binding of the PfCSP repeat by inhibitory antibodies has been shown to induce various conformations in this intrinsically disordered region (37, 39, 40, 42–44, 59, 60). Here, we show that the PbCSP repeat adopts an extended and bent conformation when recognized by inhibitory mAb 3D11. Antibody recognition of the PfCSP repeat is often mediated by aromatic cages formed by the paratope, which surround prolines, backbone atoms and aliphatic portions of side chains in the epitope. Antibody paratope residues partaking in aromatic cages often include germline-encoded residues, such as H.Trp52 from VH3-33 signature genes that are strongly recruited in the humoral response against PfCSP (38, 60, 61). Similarly, murine mAb 3D11 uses eight aromatic residues to recognize the PbCSP repeat. Germline-encoded K.Tyr32 appears to play a central role in mAb 3D11 PbCSP recognition by contacting consecutive Asn-Asp-Pro residues (PN(A/P)<u>NDP</u>(A/P)P) in the middle of the core epitope, contributing 58 Å^2^ of BSA on the Fab. These findings indicate a central role for germline-encoded aromatic residues in antibody binding of *Plasmodium* CSP repeats across species.

Our structural and biophysical data demonstrated that mAb 3D11 is cross-reactive and binds the different repeat motifs of PbCSP in nearly identical conformations. Such cross-reactivity is also exhibited by inhibitory human antibodies encoded by a variety of Ig-gene combinations when binding repeat motifs of subtle differences in PfCSP (39, 40, 43, 44, 62). Moreover, it was previously reported that human anti-PfCSP antibody affinity is often directly associated with epitope cross-reactivity (40). Given the high inhibitory potency of mAb 3D11, our findings suggest that favorable selection of cross-reactive clones during B cell maturation may have evolved as a common mechanism of the immune response in mammals against *Plasmodium* CSP.

Most residues that mediate mAb 3D11 contacts with the PbCSP repeat are germline-encoded; indeed, of nine affinity-matured residues in the HC and three in the KC, only three are involved in direct contacts with the antigen (H.Trp50, H.Val56 and K.Tyr27D). Due to the repetitive nature of the central repeat motifs, multiple antibodies bind simultaneously to one CSP protein and neighboring Fabs engage in homotypic interactions (37, 41). Our data suggest that somatic mutations of residues that partake in Fab-Fab contacts enhance homotypic interactions and indirectly improve the binding affinity of the mAb to PbCSP. In this aspect, mAb 3D11 binding to PbCSP resembles binding of some neutralizing human mAbs to PfCSP (37, 40, 41). Taken together, these findings indicate that homotypic interactions are a feature by which the mammalian immune system can robustly engage repetitive *Plasmodium* antigens with high affinity. Interestingly, recent studies have reported that Fab-Fab interactions occur in other antibody-antigen complexes, providing evidence that homotypic contacts can drive diverse biology: for example, homotypic interactions were described between two nanobodies bound to a pentameric antigen (63) and between two Rituximab antibodies when bound to B cell membrane protein CD20 (64).

Our cryoEM analysis also revealed how the PbCSP repeat, like that of PfCSP, can adopt a highly organized spiral structure upon mAb binding. Such spiral assembly of CSP was previously observed upon human mAb 311 Fab and IgG binding, which induced a PfCSP spiral with a greater diameter (27 Å) and smaller pitch (49 Å) compared to the 3D11-PbCSP complex (16 Å diameter and 51 Å pitch) (41) (**Fig. S8**). Differences in the architecture of these two complexes can be attributed to the fact that mAbs 3D11 and 311 recognize their respective antigens in distinct conformations. Because different anti-CSP inhibitory antibodies can bind the repeat region in a variety of conformations (37, 39, 43, 44, 59), it is likely that many types of CSP-antibody assemblies exist. Further studies are needed to investigate whether formation of highly organized complexes is possible on the surface of live sporozoites and how antibody-CSP interactions occur in the context of polyclonal serum. These insights will be important for our structure-function understanding of the mechanisms employed by these repeat-targeting antibodies to inhibit sporozoite development, migration and infection of hepatocytes.

## ACKNOWLEDGEMENTS

We are grateful to Dr. Samir Benlekbir for help with cryoEM data collection and for advice regarding specimen preparation. This work was supported by the CIFAR Azrieli Global Scholar program (JPJ), the Ontario Early Researcher Award program (JPJ), the Canada Research Chair program (JPJ and JLR), and the Canadian Institutes of Health Research (RP). E.T. was supported by a CIHR Canada Graduate Scholarship. A.S. was supported by an NSERC Canada Graduate Scholarship. This research was enabled in part by support provided by Compute Ontario (www.computeontario.ca) and Compute Canada (www.computecanada.ca). The ITC and BLI instruments were accessed at the Structural & Biophysical Core Facility, The Hospital for Sick Children, supported by the Canada Foundation for Innovation and Ontario Research Fund. CryoEM data was collected at the Toronto High Resolution High Throughput cryoEM facility, supported by the Canada Foundation for Innovation and Ontario Research Fund. X-ray diffraction experiments were performed at GM/CA@APS, which has been funded in whole or in part with federal funds from the National Cancer Institute (ACB-12002) and the National Institute of General Medical Sciences (AGM-12006). The Eiger 16M detector was funded by an NIH–Office of Research Infrastructure Programs High-End Instrumentation grant (1S10OD012289-01A1). This research used resources of the Advanced Photon Source, a U.S. Department of Energy (DOE) Office of Science user facility operated for the DOE Office of Science by Argonne National Laboratory under contract DE-AC02-06CH11357. X-ray diffraction experiments were also performed at the National Synchrotron Light Source II, a U.S. Department of Energy (DOE) Office of Science User Facility operated for the DOE Office of Science by Brookhaven National Laboratory under Contract No. DE-SC0012704. The Life Science Biomedical Technology Research resource is primarily supported by the National Institute of Health, National Institute of General Medical Sciences (NIGMS) through a Biomedical Technology Research Resource P41 grant (P41GM111244), and by the DOE Office of Biological and Environmental Research (KP1605010). The following reagent was obtained through BEI Resources, NIAID, NIH: Hybridoma 3D11 AntiPlasmodium berghei 44-Kilodalton Sporozoite Surface Protein (Pb44), MRA-100, contributed by Victor Nussenzweig. X-ray crystallography and cryoEM data and structures have been deposited to the Protein Data Bank and the Electron Microscopy Data Bank under PDB IDs 6X8P, 6X8Q, 6X8S, 6X8U and 6X87, and EMDB 22089, respectively.

## MATERIALS AND METHODS

### Molecular dynamics simulations

We performed all-atom molecular dynamics simulations of the following peptides: (NPNA)_5_, KQPADGNPDPNANPN, NPDPNANPNVDPNANP, (NVDPNANP)_2_NVDP, (PPPPNPND)_2_, (PPPPNAND)_2_, (PAPPNAND)_2_, and PPPPNPNDPAPPNAND as blocked monomers in water with 0.15M NaCl. Each simulation system consisted of the respective peptide with an acetylated N-terminus and amidated C-terminus solvated in a dodecahedral box with side lengths of 4.9 nm.

The systems were simulated using the program GROMACS 5.1.4 (65, 66) with the CHARMM22* (67–71) force field for the protein and the TIP3P (72) water model. All simulations were performed with periodic boundary conditions at a constant pressure and temperature of 1 bar and 300 K, respectively. The LINCS algorithm was used to constrain all bond lengths (73, 74). A cut-off of 1.4 nm was used for Lennard-Jones interactions. The Particle-Mesh Ewald algorithm (75, 76) was used to calculate long-range electrostatics interactions with a Fourier spacing of 0.12 and an interpolation order of 4. The Nosé-Hoover thermostat (77, 78) was used for temperature coupling with the peptide and solvent coupled to two temperature baths and a time constant of 0.1 ps. The Parrinello-Rahman algorithm (79) was used for pressure coupling with a time constant of 2 ps. The integration step size was 2 fs and the system coordinates were stored every 10 ps.

The simulations were performed for 300 ns for 20 independent replicas of (NPNA)_5_ and 10 independent replicas of all other sequences. The initial structures of the peptides were selected from 10 ns simulations in which extended conformations of the peptides were collapsed *in vacuo*. The first 100 ns of each trajectory were omitted as the time required for system relaxation based on the convergence analysis of the radius of gyration (Rg) shown in **Fig. S2**. This protocol resulted in a total of 4 μs of production time for the (NPNA)5 dataset and a total of 2 μs of production time for the other systems, which was used to compute equilibrium ensemble properties. The peptide snapshots were generated with VMD (80) and the plots were created with Matplotlib (81).

### 3D11 Fab production and purification

The mAb 3D11-expressing hybridoma cell line variable heavy and light chain antibody genes were sequenced (Applied Biological Materials Inc). mAb 3D11 V_K_ and V_H_ regions were cloned individually into custom pcDNA3.4 expression vectors immediately upstream of human Igκ and Igγ1-C_H_1 domains, respectively. The resulting 3D11 Fab light and heavy chain vectors were cotransfected into HEK293F cells for transient expression, and purified via KappaSelect affinity chromatography (GE Healthcare), cation exchange chromatography (MonoS, GE Healthcare), and size exclusion chromatography (Superdex 200 Increase 10/300 GL, GE Healthcare).

### Recombinant PbCSP production and purification

A construct of full-length PbCSP, the PbCSP C-terminal domain and the PbCSP αTSR domain from strain ANKA (NCBI reference sequence XP_022712148.1, residues 24-318) were designed with potential N-linked glycosylation sites mutated to glutamine and cloned into pcDNA3.4 expression vectors. The resulting constructs were transiently transfected in HEK293F cells, and purified by HisTrap FF affinity chromatography (GE Healthcare) and size exclusion chromatography (Superdex 200 Increase 10/300 GL, GE Healthcare).

### Binding kinetics by biolayer interferometry

BLI (Octet RED96, FortéBio) experiments were conducted to determine the binding kinetics of the 3D11 Fab to recombinant PbCSP. PbCSP was diluted to 10 μg/ml in kinetics buffer (PBS, pH 7.4, 0.01% [w/v] BSA, 0.002% [v/v] Tween-20) and immobilized onto Ni-NTA (NTA) biosensors (FortéBio). After a steady baseline was established, biosensors were dipped into wells containing twofold dilutions of 3D11 Fab in kinetics buffer. Tips were then immersed back into kinetics buffer for measurement of the dissociation rate. Kinetics data were analyzed using the FortéBio’s Data Analysis software 9.0, and curves were fitted to a 2:1 binding model.

### Binding thermodynamics by isothermal titration calorimetry

Calorimetric titration experiments were performed with an Auto-iTC_200_ instrument (Malvern) at 37°C. Full-length PbCSP and PbCSP-derived peptides (GenScript) were diluted in Tris-buffered saline (TBS; 20 mM Tris, pH 8.0 and 150 mM sodium chloride) and added to the calorimetric cell. Titrations were performed with 3D11 Fab in the syringe, diluted in TBS, in 15 successive injections of 2.5 μl. Full-length PbCSP was diluted to 5 μM and titrated with 3D11 Fab at 400 μM. All PbCSP-derived peptides were diluted to 20 μM and titrated with 3D11 Fab at 200-300 μM, with the exception of the NPNDx2 peptide that was diluted to 9-10 μM and titrated with 180-200 μM 3D11 Fab. Experiments were performed at least two times, and the mean and standard error of the mean are reported. The experimental data were analyzed using Origin 7.0 according to a 1:1 binding model.

### Size-exclusion chromatography-multi-angle light scattering (SEC/MALS)

Full-length PbCSP was co-complexed with a molar excess of 3D11 Fab and loaded on a Superose 6 Increase 10/300 GL (GE Healthcare) using an Agilent Technologies 1260 Infinity II HPLC coupled in-line with the following calibrated detectors: (i) MiniDawn Treos MALS detector (Wyatt); (ii) Quasielastic light scattering (QELS) detector (Wyatt); and (iii) Optilab T-reX refractive index (RI) detector (Wyatt). Data processing was performed using the ASTRA software (Wyatt).

### Crystallization and structure determination

Purified 3D11 Fab was mixed in a 1:5 molar ratio with each PbCSP peptide. The 3D11 Fab/PAPP complex was concentrated to 5 mg/mL and mixed in a 1:1 ratio with 20% (w/v) PEG 3350, 0.15 M malic acid pH 7. Crystals appeared after ~1 d and were cryoprotected in 15% (v/v) ethylene glycol before being flash-frozen in liquid nitrogen. The 3D11 Fab/NAND complex was concentrated to 5 mg/mL and mixed in a 1:1 ratio with 20% (w/v) PEG 3350, 0.2 M di-sodium tartrate. Crystals appeared after ~3 d and were cryoprotected in 15% (v/v) ethylene glycol before being flash-frozen in liquid nitrogen. The 3D11 Fab/NPND complex was concentrated to 2.1 mg/mL and mixed in a 1:1 ratio with 25% (w/v) PEG 3350, 0.2 M lithium sulfate, 0.1 M Tris pH 8.5. Crystals appeared after ~12 d and were cryoprotected in 20% (v/v) ethylene glycol before being flash-frozen in liquid nitrogen. The 3D11 Fab/Mixed complex was concentrated to 5 mg/mL and mixed in a 1:1 ratio with 25.5% (w/v) PEG 4000, 15% (v/v) glycerol, 0.17 M ammonium acetate, 0.085 M sodium citrate pH 5.6. Crystals appeared after ~ 1 d and were cryoprotected in 20% (v/v) glycerol before being flash-frozen in liquid nitrogen.

Data were collected at the 23-ID-D or 23-ID-B beamline at the Argonne National Laboratory Advanced Photon Source, or at the 17-ID-1 beamline at the National Synchrotron Light Source II. All datasets were processed and scaled using XDS (82). The structures were determined by molecular replacement using Phaser (83). Refinement of the structures was performed using phenix.refine (84) and iterations of refinement using Coot (85). Access to all software was supported through SBGrid (86).

### CryoEM data collection and image processing

The PbCSP/3D11 complex was concentrated to 3 mg/mL and incubated briefly with 0.01% (w/v) n-Dodecyl β-D-maltopyranoside. 3 μl of the sample was deposited on holey gold grids prepared in-house (87), which were glow discharged in air for 15 s before use. Sample was blotted for 12.5 s, and subsequently plunge-frozen in a mixture of liquid ethane and propane (88) using a modified FEI Vitrobot (maintained at 4°C and 100% humidity). Data collection was performed on a Thermo Fisher Scientific Titan Krios G3 operated at 300 kV with a Falcon 3EC camera automated with the EPU software. A nominal magnification of 75,000× (calibrated pixel size of 1.06 Å) and defocus range between 1.6 and 2.2 μm were used for data collection. Exposures were fractionated as movies of 30 frames with total exposure of 42.7 electrons/Å^2^. A total of 2,080 raw movies were obtained.

Image processing was carried out in cryoSPARC v2 (89). Initial specimen movement correction, exposure weighting, and CTF parameters estimation were done using patch-based algorithms. Manual particle selection was performed on 30 micrographs to create templates for the template-based picking. 669,223 particle images were selected by template picking and individual particle images were corrected for beam-induced motion with the local motion algorithm (90). *Ab-initio* structure determination revealed that most particles present in the dataset correspond to the 3D11 Fab-PbCSP complex, with a minor population of particles corresponding to unbound 3D11 Fab. After several rounds of heterogeneous refinement, 165,747 particle images were selected for non-uniform refinement with no symmetry applied, which resulted in a 3.2 Å resolution map of the 3D11 Fab-PbCSP complex estimated from the gold-standard Fourier shell correlation (FSC) criterion.

### Model building

To create a starting model of the 3D11 Fab-PbCSP complex, seven copies of the Fab3D11/PbCSP-peptide crystal structure were manually docked into the 3D11 Fab-PbCSP cryoEM map using UCSF Chimera (91), followed by manual building using Coot (85). All models were refined using the phenix.real_space_refine (84) with secondary structure and geometry restraints. The final models were evaluated by MolProbity (92).

## Supplementary Figures Legend

**Figure S1.**
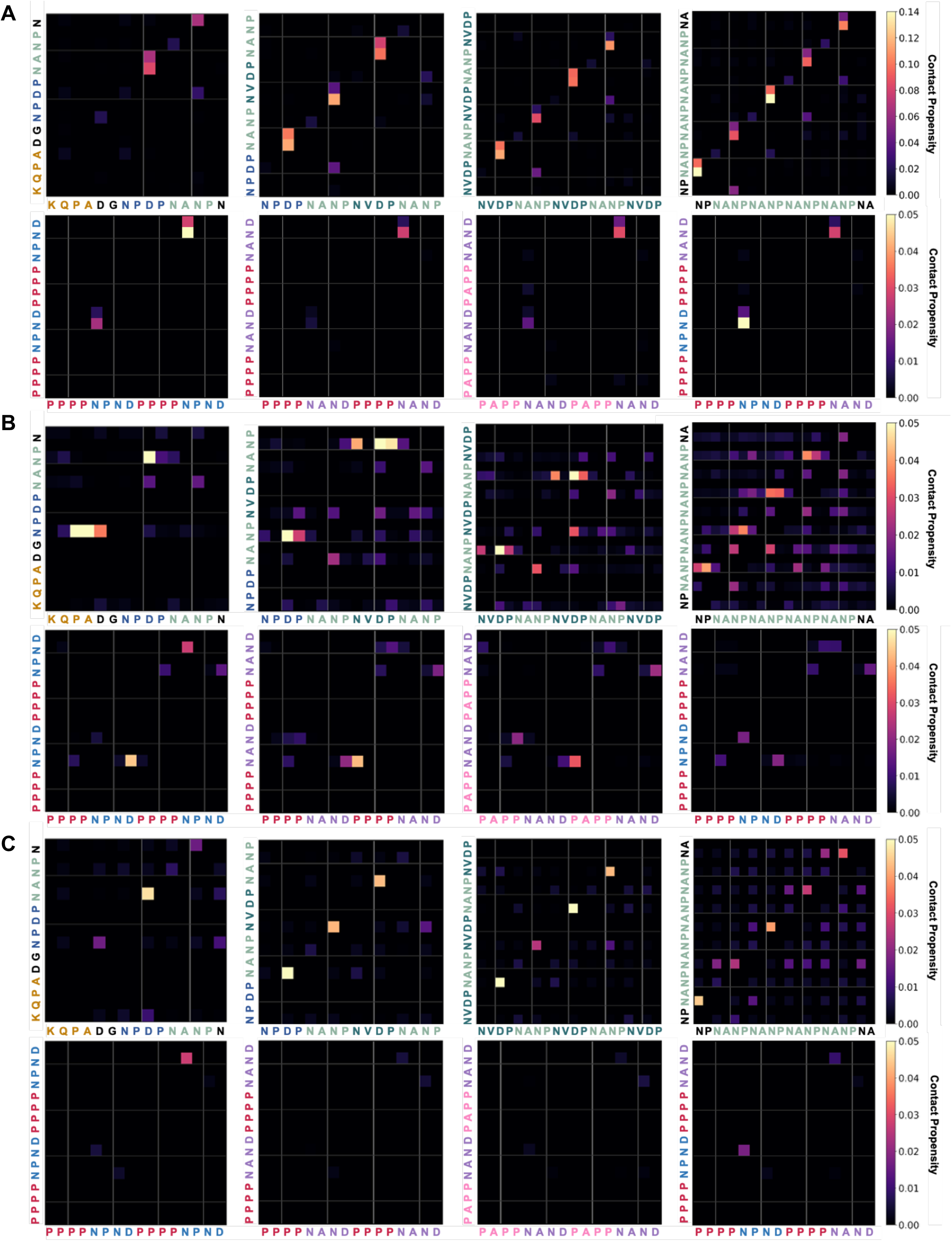
Ensemble-averaged hydrogen-bonding propensities for PfCSP- and PbCSP-derived peptides. **(A)** The propensity for hydrogen bonds between the NH groups of the backbone (y-axis) and CO groups of the side chains (x-axis) is indicated by the color scale on the right. **(B)** The propensity for hydrogen bonds between the NH groups of the side chains (y-axis) and CO groups of the backbone (x-axis) is indicated by the color scale on the right. **(C)** The propensity for hydrogen bonds between the NH groups of the side chains (y-axis) and CO groups of the side chains (x-axis) is indicated by the color scale on the right.

**Figure S2.**
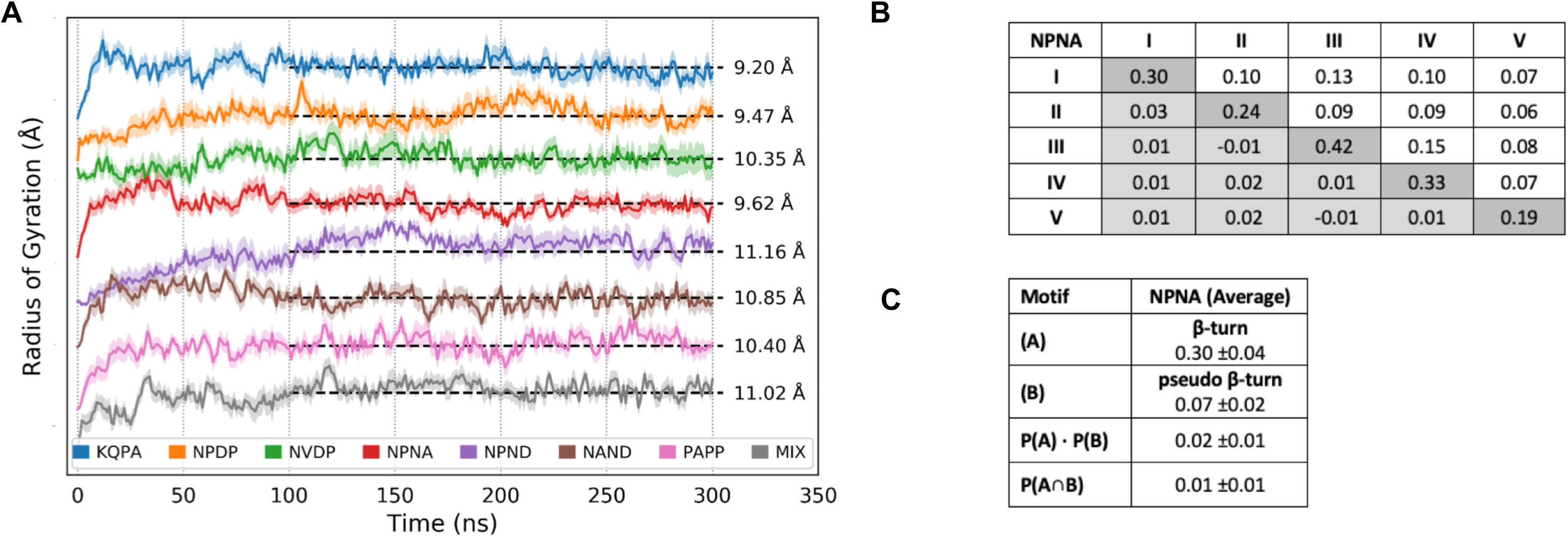
Experimental details of MD simulations. **(A)** Time evolution of the radius of gyration of the different peptides from MD simulations. Shading represents the standard error of the mean computed from the different simulation repeats (see Methods). The simulations are statistically converged after 100 ns. The average radius of gyration for each peptide is reported on the right. The individual peptides have been shifted on the y-axis to allow for ease of visualization. **(B)** The simulated propensity of having a specific hydrogen bond is shown on the diagonal, with the propensity of two specific hydrogen bonds (P(A⋂B)) shown above the diagonal. Each individual NPNA motif is labelled from I-V. The difference of the simulated values from the calculated values (P(A)·P(B)) is shown below the diagonal. These differences are all well below the average standard error of mean of 0.04, confirming that the individual motifs are uncorrelated. **(C)** A mathematical example showing that the β-turn and pseudo β-turn propensities are independent and uncorrelated (P(A⋂B) = P(A)·P(B)). This holds true for all the simulated peptides.

**Figure S3.**
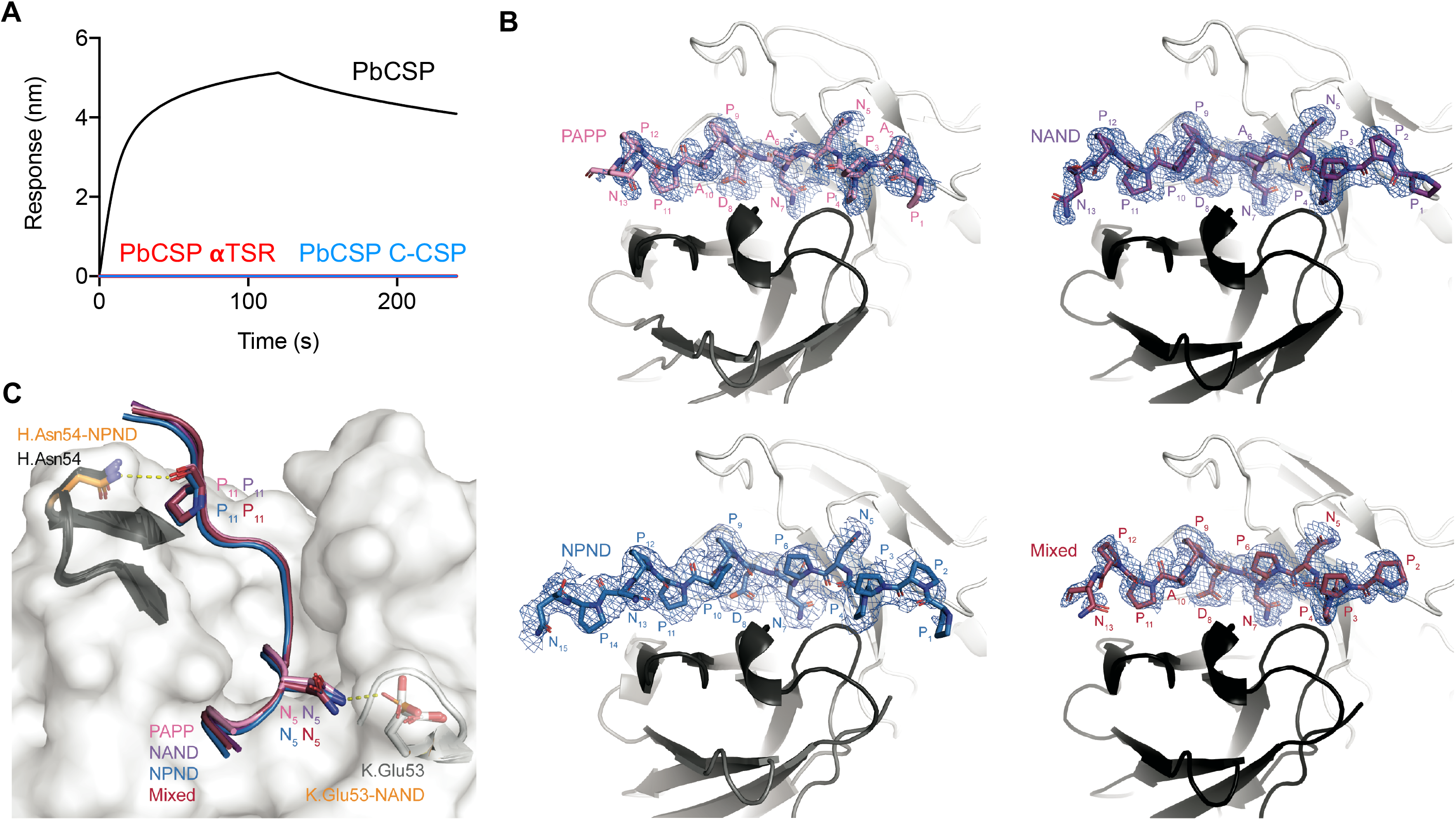
Experimental details of mAb 3D11 binding. **(A)** 3D11 Fab binding to peptides representative of the PbCSP aTSR domain (residue 263-318; red) and the full C-terminal domain (residues 202-318; C-CSP; blue). Full-length PbCSP (residue 24-318) was used as a positive control (black). **(B)** Composite omit map electron density contoured at 1.0 sigma (blue mesh) around PbCSP peptides PAPP, NAND, NPND and Mixed in complex with the 3D11 Fab. **(C)** Slight differences in H-bonding at the N- and C-terminal ends of the PbCSP repeat peptides when bound to the 3D11 Fab in the crystal structures. PbCSP peptides are colored as in **Fig. 3**. Antibody residues partaking in H-bonds are colored orange. mAb 3D11 HCDR2 is colored in black and KCDR2 in white.

**Figure S4.**
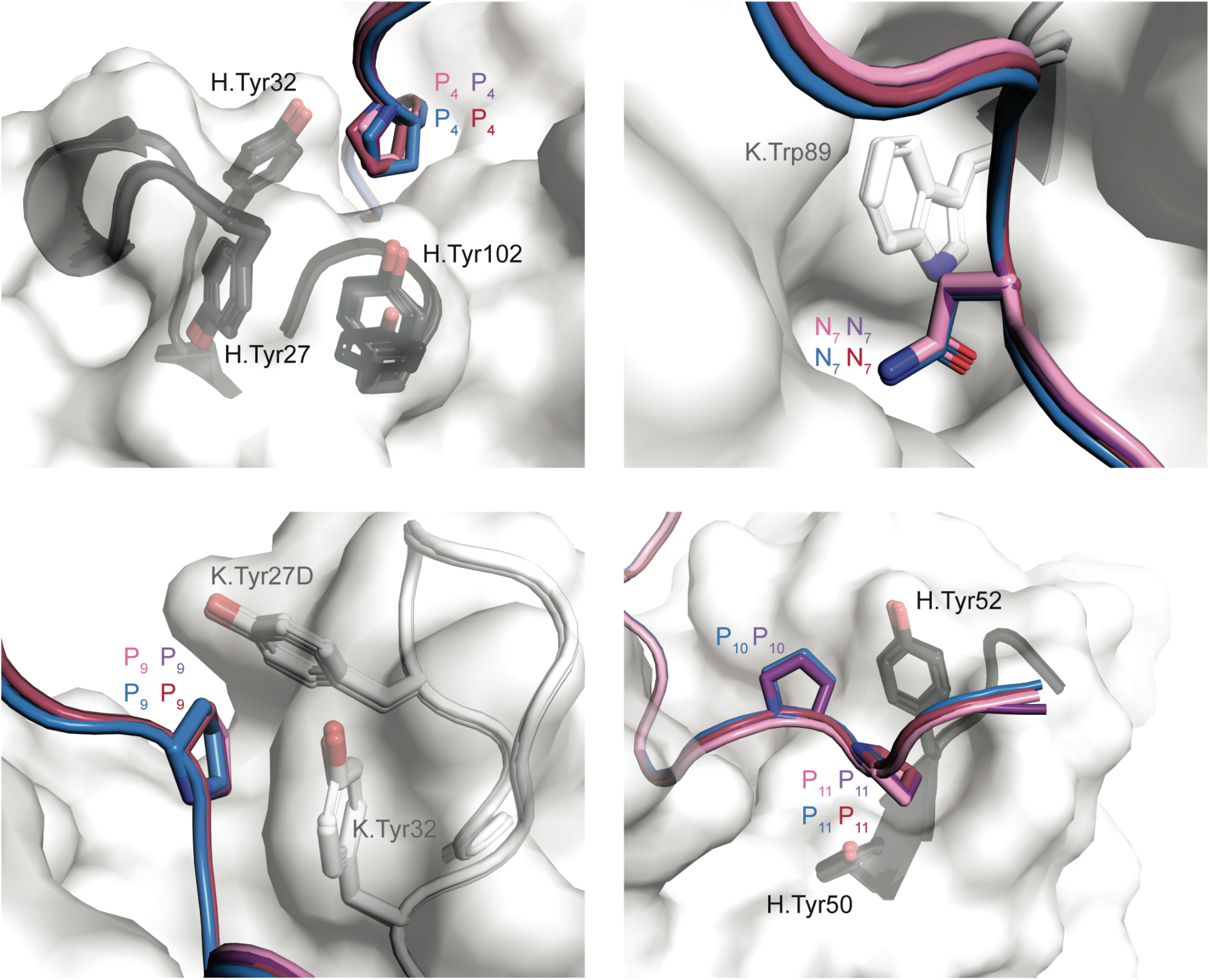
Interactions between mAb 3D11 aromatic side chains and PbCSP peptides. Interactions formed between PbCSP peptide residues and aromatic side chains of the 3D11 Fab HCDR (black) and KCDR (white). PbCSP peptides are colored as in **Fig. 3.**

**Figure S5.**
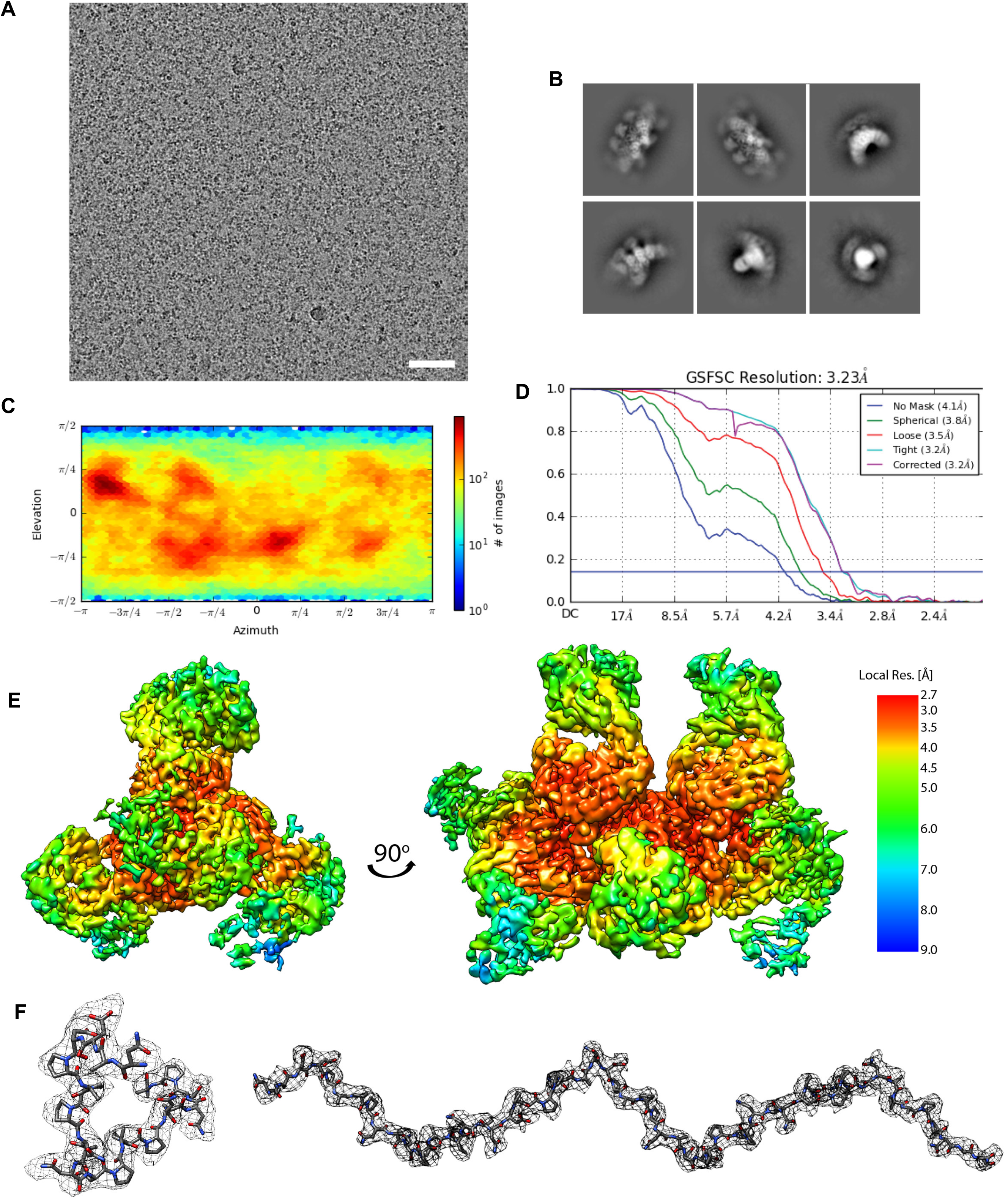
CryoEM analysis of the 3D11 Fab-PbCSP complex. **(A)** A representative cryoEM micrograph. Scale bar, 50 nm. **(B)** Selected 2D class averages of the 3D11 Fab-PbCSP complex. **(C)** Particle orientation distribution plot. **(D)** Fourier shell correlation curve from the final 3D non-uniform refinement of the 3D11 Fab-PbCSP complex in cryoSPARC v2. **(E)** Local resolution (Å) plotted on the surface of the cryoEM map. (**F**) CryoEM density of PbCSP (grey mesh) with the model built shown as sticks (black carbons).

**Figure S6.**
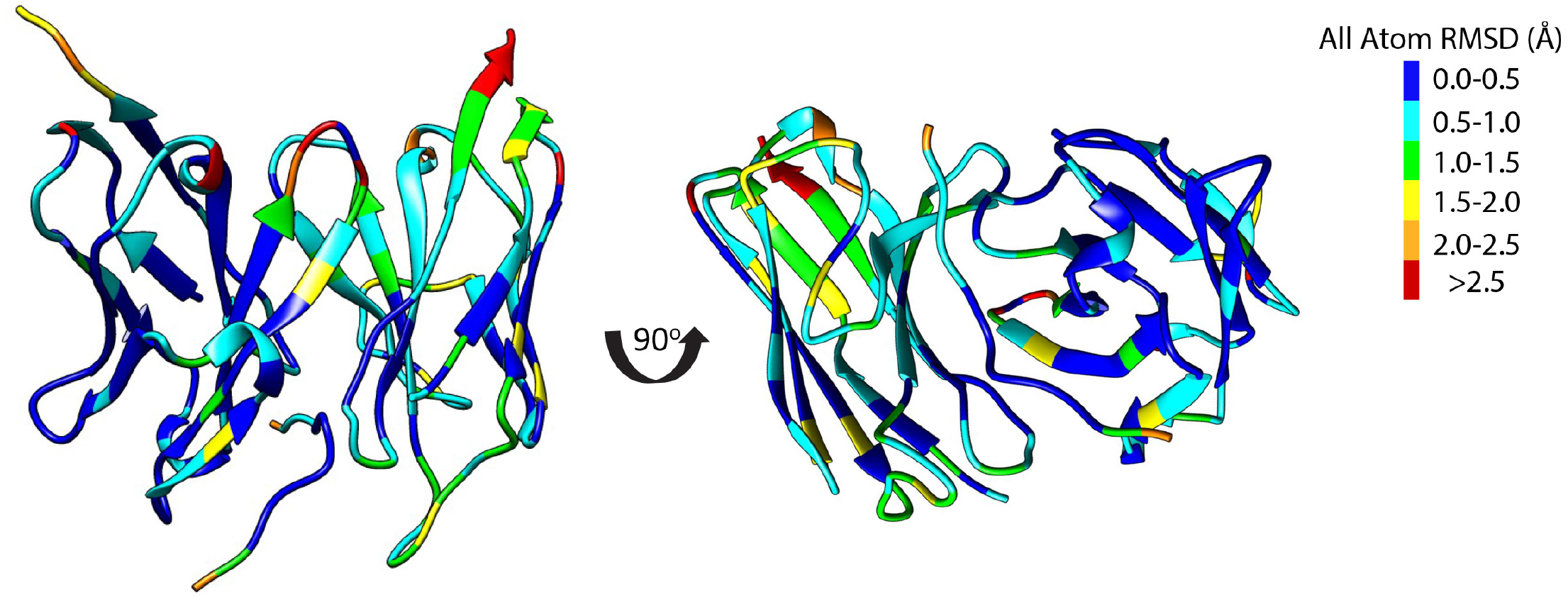
Comparison between 3D11 Fab-PbCSP cryoEM structure and 3D11 Fab-NPND peptide crystal structure. Color representation of all-atom RMSD of 3D11 Fab variable region and PbCSP core epitope between the Fab-PbCSP cryoEM structure and the 3D11-NPND peptide crystal structure. RMSD values were calculated using UCSF Chimera (91) and plotted by color on the secondary structure of the 3D11-NPND peptide crystal structure.

**Figure S7.**
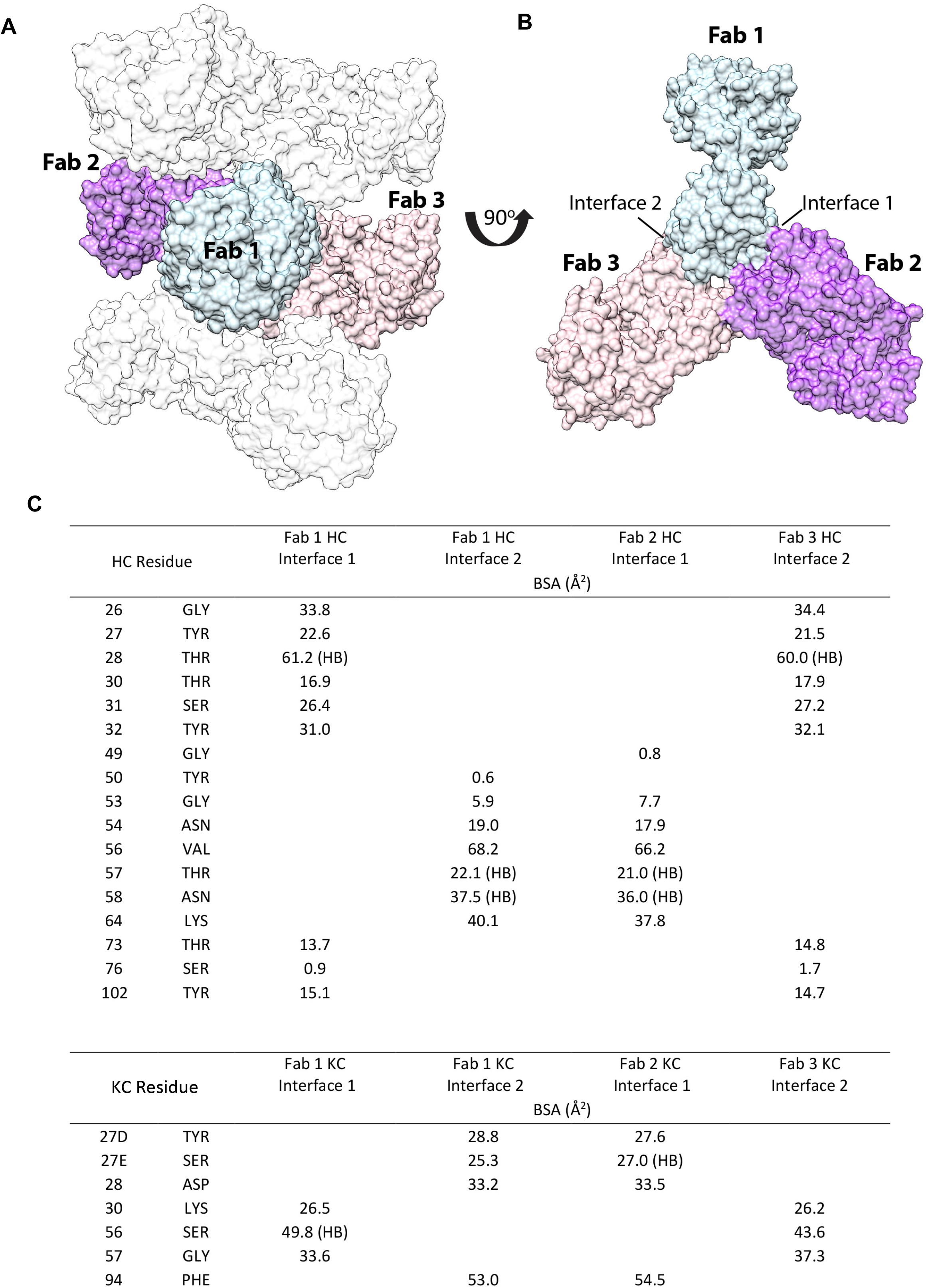
Homotypic contacts between 3D11 Fabs in the 3D11 Fab-PbCSP cryoEM structure. (**A**) and (**B**) main interaction interfaces between adjacent 3D11 Fabs in the 3D11 Fab-PbCSP cryoEM structure. (**C**) Table of contacts between 3D11 Fabs. HB: hydrogen bond (3.8 Å cut-off).

**Figure S8.**
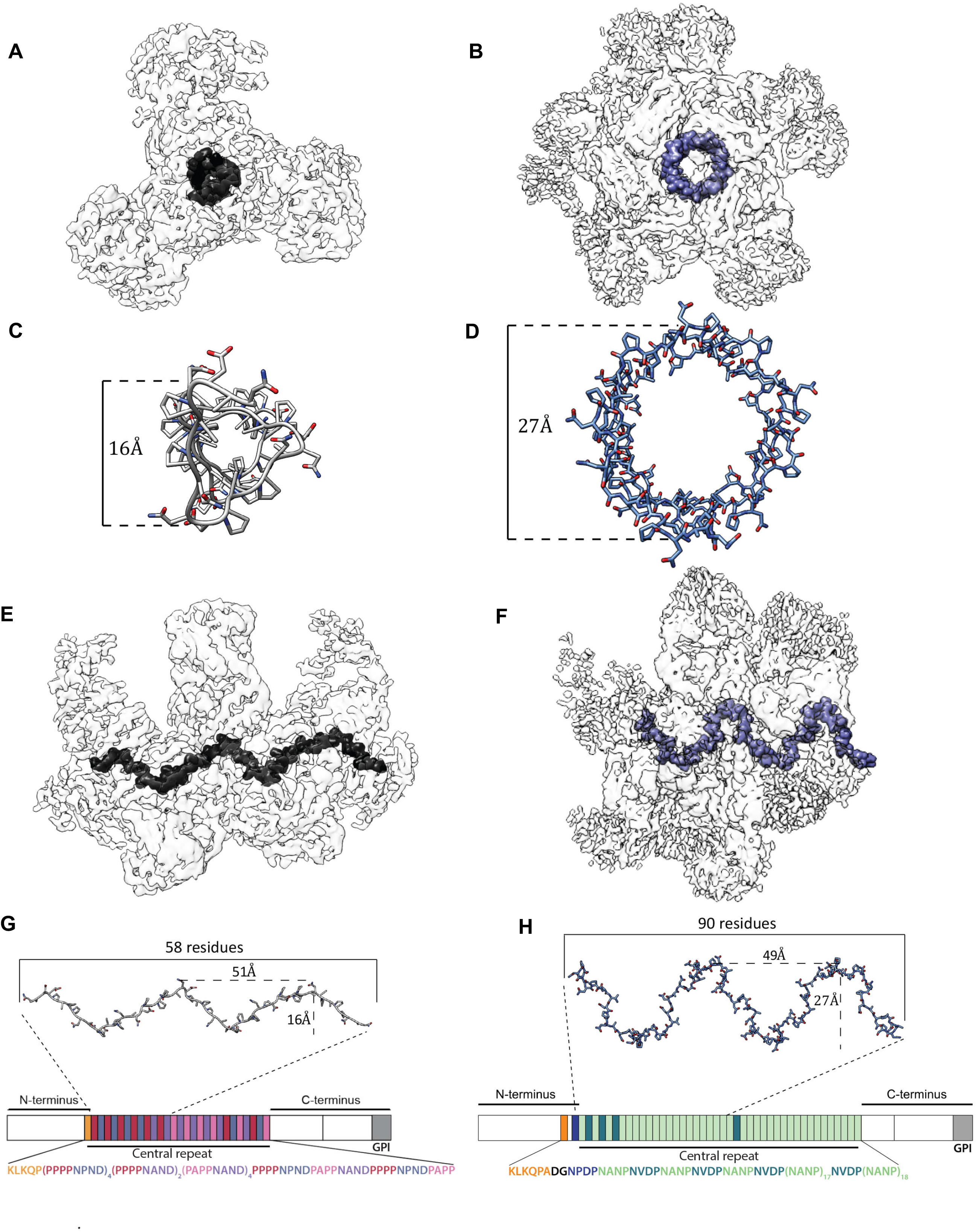
Comparison between cryoEM structures of 3D11 Fab-PbCSP and 311 Fab-PfCSP. (PDB ID: 6MB3) (41). CryoEM maps of the 3D11 Fab-PbCSP ((**A**) and (**E**)) and 311 Fab-PfCSP ((**B**) and (**F**)) complexes are shown as a transparent light gray surface with the CSP density highlighted in black for PbCSP and in blue for PfCSP. The CSP models built into the cryoEM maps are shown as grey (for PbCSP; (**C**) and (**F**)) or blue (for PbCSP; (**D**) and (**H**)) sticks and aligned to the schematic representations of their respective protein sequences.

**Table S1.**
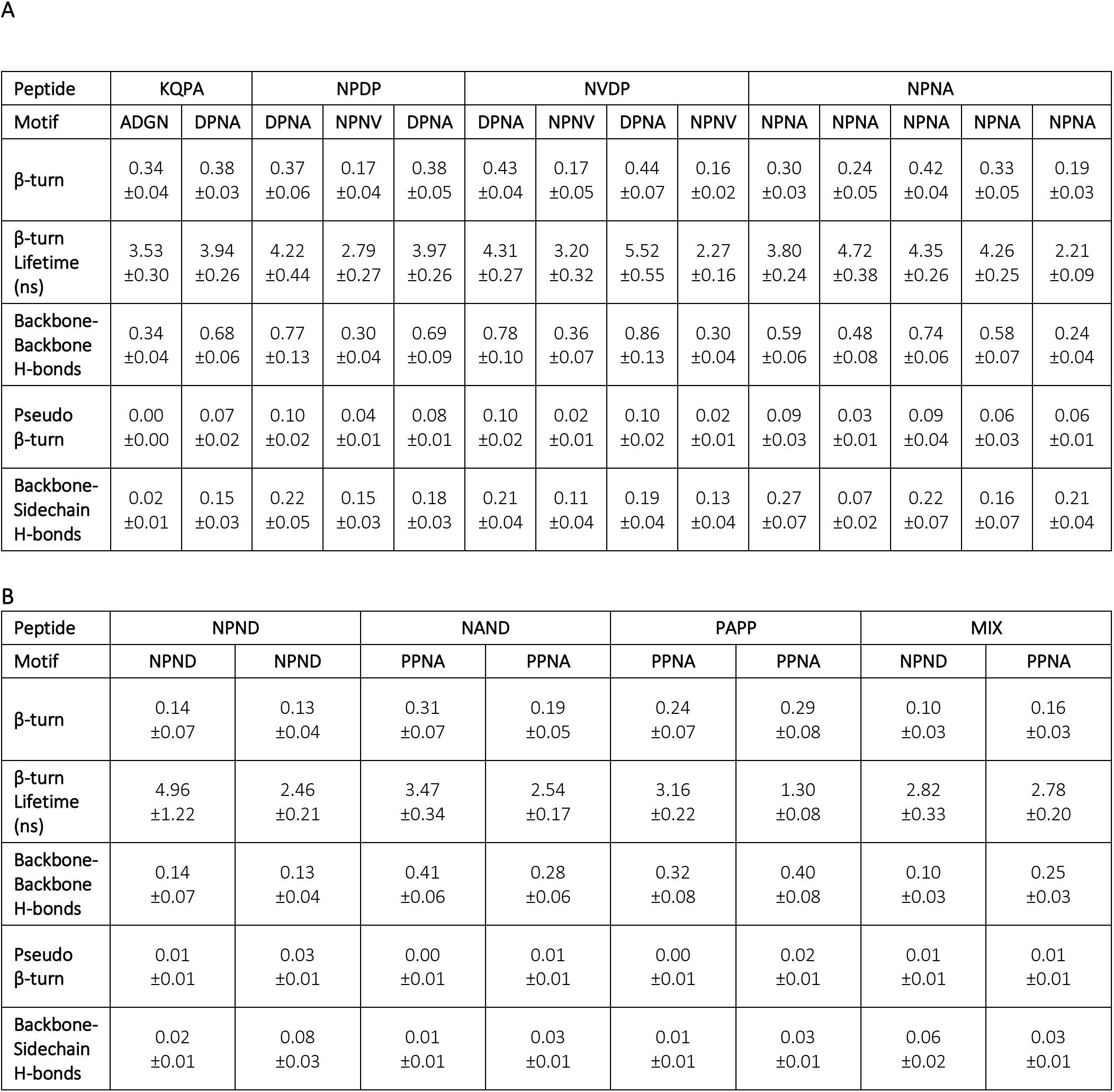
Hydrogen bonding propensities from simulations of peptides in solution. **(A)** Hydrogen-bonding propensity for each simulated motif and lifetime of each β-turn for the four PfCSP-derived peptides. **(B)** Hydrogen-bonding propensity for each simulated motif and lifetime of each β-turn for the four PbCSP-derived peptides.

| Peptide | KQPA |  | NPDP |  |  | NVDP |  |  |  | NPNA |  |  |  |  |
| --- | --- | --- | --- | --- | --- | --- | --- | --- | --- | --- | --- | --- | --- | --- |
| Motif | ADGN | DPNA | DPNA | NPNV | DPNA | DPNA | NPNV | DPNA | NPNV | NPNA | NPNA | NPNA | NPNA | NPNA |
| β-turn | 0.34<br>$\pm 0.04$ | 0.38<br>$\pm 0.03$ | 0.37<br>$\pm 0.06$ | 0.17<br>$\pm 0.04$ | 0.38<br>$\pm 0.05$ | 0.43<br>$\pm 0.04$ | 0.17<br>$\pm 0.05$ | 0.44<br>$\pm 0.07$ | 0.16<br>$\pm 0.02$ | 0.30<br>$\pm 0.03$ | 0.24<br>$\pm 0.05$ | 0.42<br>$\pm 0.04$ | 0.33<br>$\pm 0.05$ | 0.19<br>$\pm 0.03$ |
| β-turn<br>Lifetime<br>(ns) | 3.53<br>$\pm 0.30$ | 3.94<br>$\pm 0.26$ | 4.22<br>$\pm 0.44$ | 2.79<br>$\pm 0.27$ | 3.97<br>$\pm 0.26$ | 4.31<br>$\pm 0.27$ | 3.20<br>$\pm 0.32$ | 5.52<br>$\pm 0.55$ | 2.27<br>$\pm 0.16$ | 3.80<br>$\pm 0.24$ | 4.72<br>$\pm 0.38$ | 4.35<br>$\pm 0.26$ | 4.26<br>$\pm 0.25$ | 2.21<br>$\pm 0.09$ |
| Backbone-Backbone<br>H-bonds | 0.34<br>$\pm 0.04$ | 0.68<br>$\pm 0.06$ | 0.77<br>$\pm 0.13$ | 0.30<br>$\pm 0.04$ | 0.69<br>$\pm 0.09$ | 0.78<br>$\pm 0.10$ | 0.36<br>$\pm 0.07$ | 0.86<br>$\pm 0.13$ | 0.30<br>$\pm 0.04$ | 0.59<br>$\pm 0.06$ | 0.48<br>$\pm 0.08$ | 0.74<br>$\pm 0.06$ | 0.58<br>$\pm 0.07$ | 0.24<br>$\pm 0.04$ |
| Pseudo<br>β-turn | 0.00<br>$\pm 0.00$ | 0.07<br>$\pm 0.02$ | 0.10<br>$\pm 0.02$ | 0.04<br>$\pm 0.01$ | 0.08<br>$\pm 0.01$ | 0.10<br>$\pm 0.02$ | 0.02<br>$\pm 0.01$ | 0.10<br>$\pm 0.02$ | 0.02<br>$\pm 0.01$ | 0.09<br>$\pm 0.03$ | 0.03<br>$\pm 0.01$ | 0.09<br>$\pm 0.04$ | 0.06<br>$\pm 0.03$ | 0.06<br>$\pm 0.01$ |
| Backbone-Sidechain<br>H-bonds | 0.02<br>$\pm 0.01$ | 0.15<br>$\pm 0.03$ | 0.22<br>$\pm 0.05$ | 0.15<br>$\pm 0.03$ | 0.18<br>$\pm 0.03$ | 0.21<br>$\pm 0.04$ | 0.11<br>$\pm 0.04$ | 0.19<br>$\pm 0.04$ | 0.13<br>$\pm 0.04$ | 0.27<br>$\pm 0.07$ | 0.07<br>$\pm 0.02$ | 0.22<br>$\pm 0.07$ | 0.16<br>$\pm 0.07$ | 0.21<br>$\pm 0.04$ |

B
| Peptide | NPND |  | NAND |  | PAPP |  | MIX |  |
| --- | --- | --- | --- | --- | --- | --- | --- | --- |
| Motif | NPND | NPND | PPNA | PPNA | PPNA | PPNA | NPND | PPNA |
| β-turn | 0.14<br>$\pm 0.07$ | 0.13<br>$\pm 0.04$ | 0.31<br>$\pm 0.07$ | 0.19<br>$\pm 0.05$ | 0.24<br>$\pm 0.07$ | 0.29<br>$\pm 0.08$ | 0.10<br>$\pm 0.03$ | 0.16<br>$\pm 0.03$ |
| β-turn<br>Lifetime<br>(ns) | 4.96<br>$\pm 1.22$ | 2.46<br>$\pm 0.21$ | 3.47<br>$\pm 0.34$ | 2.54<br>$\pm 0.17$ | 3.16<br>$\pm 0.22$ | 1.30<br>$\pm 0.08$ | 2.82<br>$\pm 0.33$ | 2.78<br>$\pm 0.20$ |
| Backbone-Backbone<br>H-bonds | 0.14<br>$\pm 0.07$ | 0.13<br>$\pm 0.04$ | 0.41<br>$\pm 0.06$ | 0.28<br>$\pm 0.06$ | 0.32<br>$\pm 0.08$ | 0.40<br>$\pm 0.08$ | 0.10<br>$\pm 0.03$ | 0.25<br>$\pm 0.03$ |
| Pseudo<br>β-turn | 0.01<br>$\pm 0.01$ | 0.03<br>$\pm 0.01$ | 0.00<br>$\pm 0.01$ | 0.01<br>$\pm 0.01$ | 0.00<br>$\pm 0.01$ | 0.02<br>$\pm 0.01$ | 0.01<br>$\pm 0.01$ | 0.01<br>$\pm 0.01$ |
| Backbone-Sidechain<br>H-bonds | 0.02<br>$\pm 0.01$ | 0.08<br>$\pm 0.03$ | 0.01<br>$\pm 0.01$ | 0.03<br>$\pm 0.01$ | 0.01<br>$\pm 0.01$ | 0.03<br>$\pm 0.01$ | 0.06<br>$\pm 0.02$ | 0.03<br>$\pm 0.01$ |

**Table S2.**
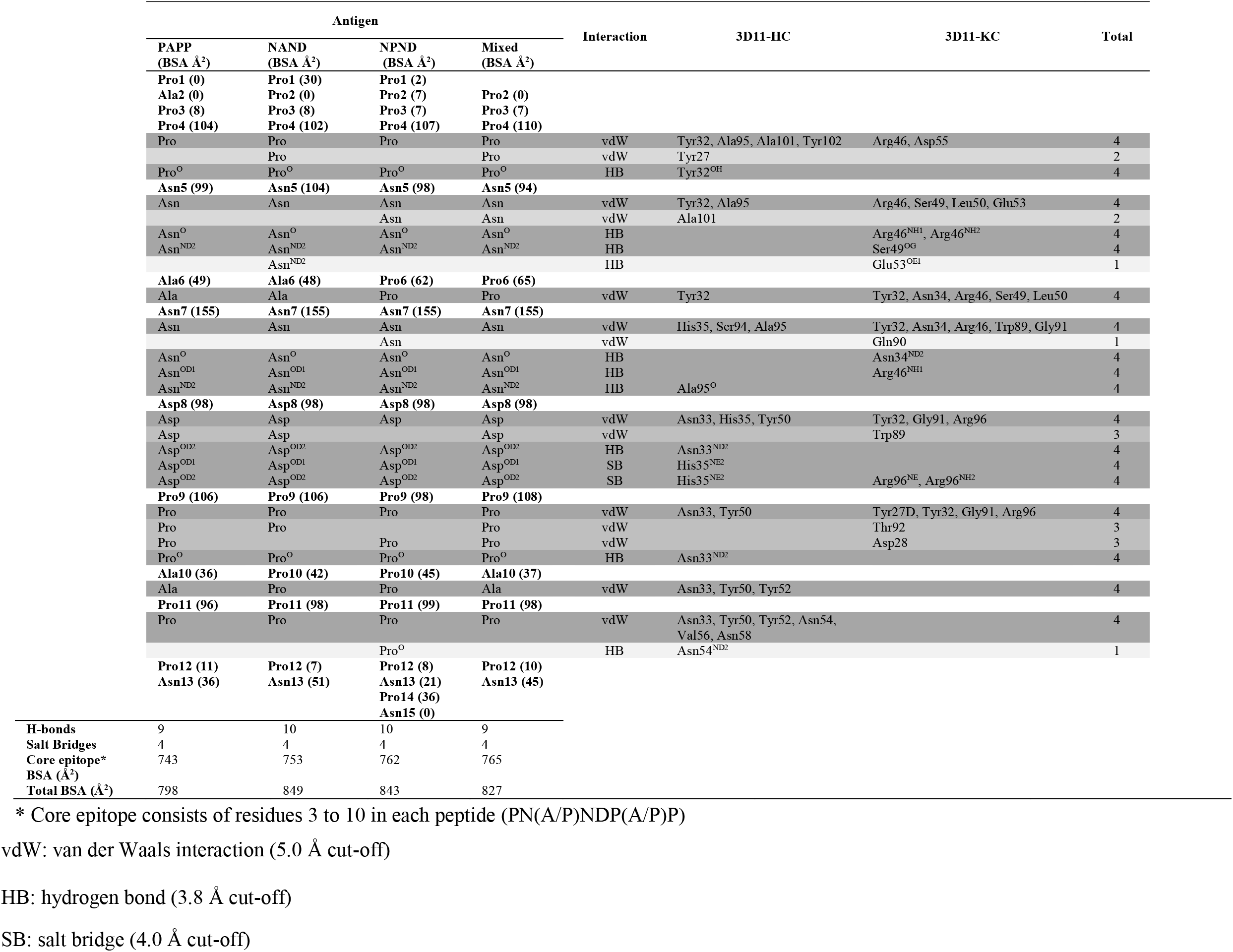
Table of contacts between 3D11 Fab and PbCSP peptides. *Rows are shaded according to the number of times interactions are observed between all four crystal structures, summed in the final column.

**Table S3.**
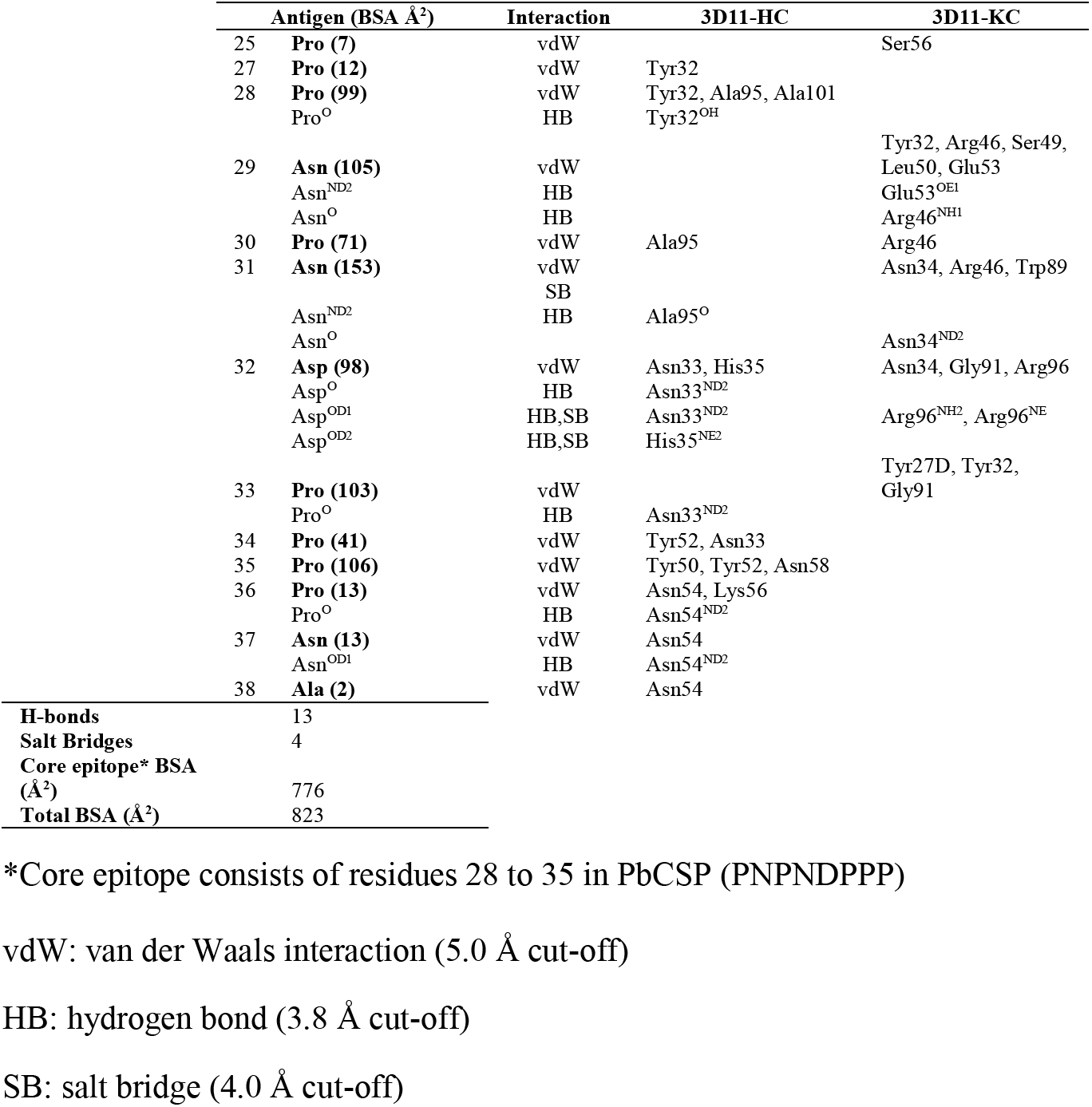
Table of contacts between one of the 3D11 Fabs and PbCSP 622 in cryoEM structure.

